# Scalable methylome-transcriptome profiling resolves clonal organization in human hematopoiesis

**DOI:** 10.64898/2026.10.08.757494

**Authors:** Jikai Shao, Mengyang Chen, Yan Liu, Jinnan Sun, Guangxin Zhang, Zheng-Yi Yan, Hongying Lin, Miaomiao Ren, Shanshan Pei, Jun Peng, Yuyan Gong, Shou-Wen Wang

## Abstract

Understanding human tissue maintenance and remodeling requires linking clonal ancestry to molecular state, yet scalable joint measurement remains challenging. Here, we introduce Droplet-MT and an integrated computational workflow for joint transcriptome and methylome profiling at ∼10,000 cells per library. Methylation epimutations enable mutation-independent clonal reconstruction from high-coverage methylomes, while reference-based label transfer assigns cells to established clones at ∼0.2% CpG coverage, reducing sequencing cost. Clone-aggregated somatic variants provide genetic annotation without additional single-cell genotyping. Across ∼21,000 paired profiles from healthy blood and longitudinal acute myeloid leukemia, we resolved small and expanded clones with heterogeneous lineage output. In leukemia, the framework distinguished changes in malignant clonal abundance, state remodeling within persistent clones, and recovery of healthy polyclonal hematopoiesis. Malignant monocytes remained transcriptionally and epigenetically distinct from healthy and recovery-associated cells despite treatment-associated remodeling. Droplet-MT enables scalable analysis of human tissues by linking clonal organization with molecular state and retrospective genetic annotation.

## Introduction

Human tissues are maintained by cellular lineages whose contributions and molecular states change during aging, regeneration and disease^1–3^. Similar cellular phenotypes can arise from distinct clones, whereas a single clone can generate descendants with different identities and functions. Consequently, a shift in tissue composition can reflect expansion of selected clones, altered lineage output within existing clones, or molecular remodeling of their descendants. Distinguishing these processes requires ancestry information alongside molecular-state measurements. The transcriptome captures current gene-expression programs, whereas DNA methylation can retain more persistent regulatory information that is not evident from RNA alone^4,5^. Integrating these complementary molecular readouts with clonal ancestry therefore provides a framework for understanding how cellular diversity is maintained and reorganized in health and disease.

Retrospective lineage tracing in humans relies on naturally occurring variants, but available genetic markers impose different constraints^2^. Nuclear somatic mutations are stable and distributed genome-wide, yet their extremely low mutation rate makes de novo discovery by single-cell whole-genome sequencing expensive and low-throughput^6–8^; colony-based strategies improve mutation detection but lose direct linkage to the original single-cell phenotype^9,10^. Mitochondrial variants are more readily captured, but heteroplasmy, intracellular genetic drift and stochastic segregation can alter their representation among descendants, complicating inference of clonal relationships^2,11–13^. Copy-number variations (CNVs) can distinguish clones in many cancers but offer limited resolution in normal tissues, where such alterations are often absent^14–16^. Targeted mutation genotyping coupled with RNA profiling can efficiently link selected variants to cellular states^15,17–20^, but its utility is similarly greatest in cancer and is limited in normal tissues lacking informative mutations. A new framework is therefore needed for scalable clonal reconstruction without predefined genetic variants, while retaining the ability to link ancestry to genotype and molecular phenotype.

DNA methylation offers a promising alternative for lineage tracing in humans. Stochastic epimutations acquired across millions of CpG sites can be inherited by descendant cells, providing orders of magnitude more candidate lineage marks per cell division than nuclear sequence mutations^21–26^. Targeted approaches such as EPI-Clone enable high-throughput lineage tracing in blood from approximately 500 predefined CpGs; however, their performance for resolving small, non-expanded lineages remains limited^21,27^. We previously developed MethylTree to infer ancestry from genome-wide epimutations while accounting for cell-state-associated methylation variation; at approximately 5% CpG coverage per cell, its reconstructions showed nearly 100% concordance with ground-truth lineage barcodes in hematopoietic models^28^. Because it does not rely on predefined mutations, this genome-wide strategy is particularly well suited to resolving polyclonal organization in healthy tissues. At the same time, the methylome reports epigenetic state and the paired transcriptome provides a complementary functional readout, linking lineage history to molecular phenotype in the same cell.

Realizing this potential at scale, however, presents three challenges. First, existing joint methylome–transcriptome methods remain substantially lower-throughput and more expensive than droplet-based single-cell RNA sequencing^25,29–31^. Second, de novo epimutation-based reconstruction benefits from an average CpG coverage of approximately 3%^28^, making discovery-depth profiling costly in large cohorts or longitudinal studies. Third, interpreting clonal behavior, particularly in cancer, requires linking ancestry and molecular state to genetic alterations^15^. Although genome-wide methylome libraries retain sequence information that can be interrogated after data collection, sparse coverage and library construction errors limit reliable cell-level genotyping.

Here, we introduce Droplet-MT and an integrated computational workflow for scalable analysis of clonal ancestry, molecular state and somatic alterations. The droplet platform generates approximately 10,000 paired transcriptome and genome-wide methylome profiles per library with sufficient CpG coverage for de novo MethylTree reconstruction. Once high-depth cells establish a clonal reference, label transfer assigns cells from represented lineages using ultra-shallow methylomes, reducing the sequencing burden of larger or longitudinal studies. Finally, aggregating mutation evidence across methylation-defined clones links inferred lineages to somatic alterations without requiring successful genotyping of every individual cell. Across ∼21,000 paired high-quality profiles from healthy blood and longitudinal acute myeloid leukemia (AML), Droplet-MT resolves heterogeneous lineage output, malignant clonal persistence and clonally diverse recovery-associated hematopoiesis, while linking these processes to transcriptional and epigenetic states.

## Results

### Droplet-MT combines scalable paired profiling with genome-wide CpG coverage

We developed Droplet-MT, a droplet-based platform, to jointly measure gene expression and genome-wide DNA methylation in single cells at high throughput. The workflow accommodates permeabilized whole cells or isolated nuclei (Fig. 1a). Following chromatin relaxation and in situ DNA tagmentation, individual cells or nuclei are encapsulated with gel beads carrying matched DNA- and RNA-capture identifiers. These identifiers link the two modalities to the same cellular input while allowing their libraries to be processed separately. Genomic DNA undergoes enzymatic C-to-T conversion for methylation readout, whereas RNA-derived cDNA is retained for transcriptome profiling. A companion computational workflow connects the paired measurements to cell-state analysis, MethylTree reconstruction and clone-resolved downstream analyses (Extended Data Fig. 1a,b).

**Fig. 1.**
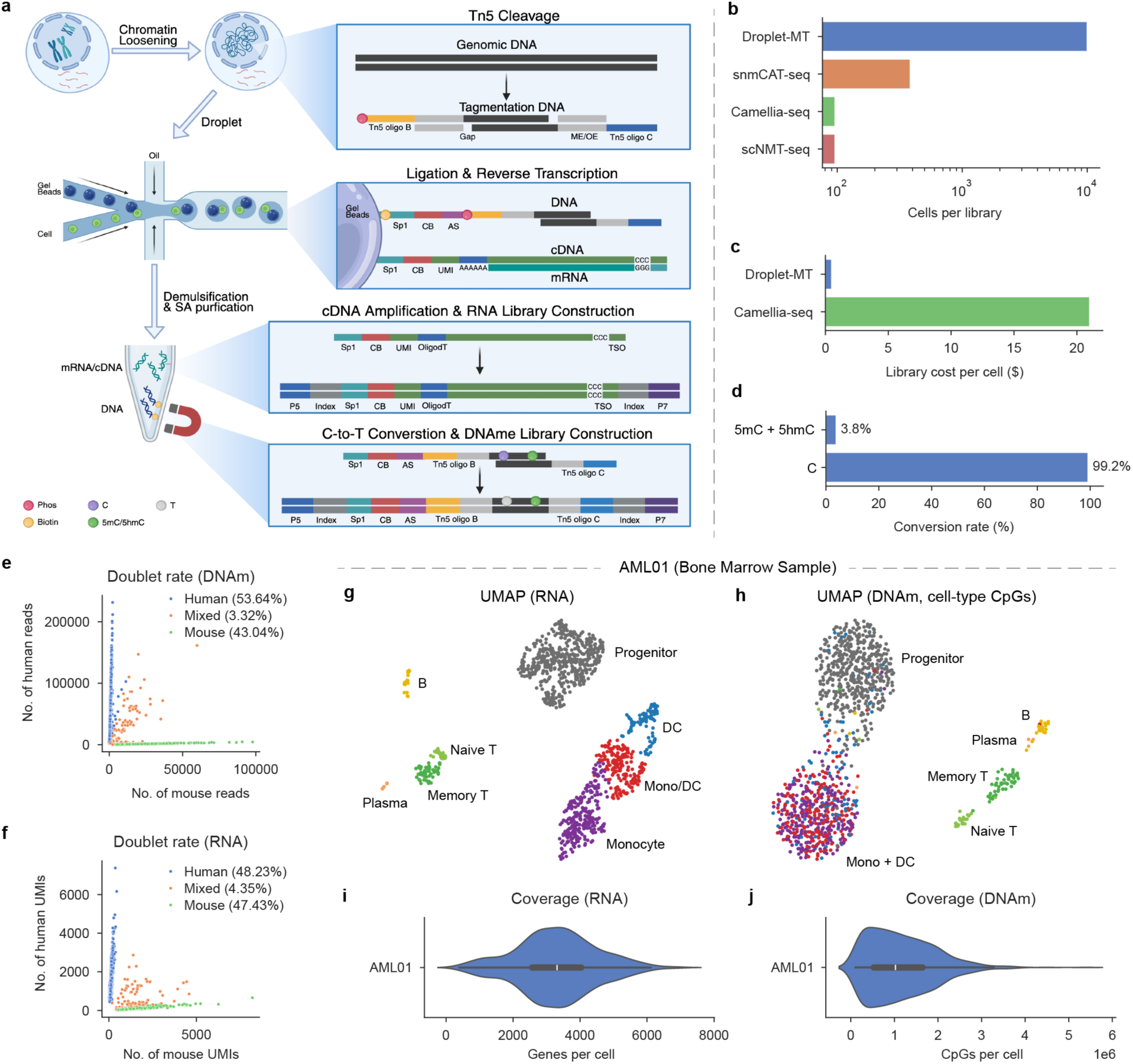
Droplet-MT enables high-throughput joint single-cell transcriptome and DNA methylome profiling. a, Schematic of the Droplet-MT workflow. Permeabilized cells are tagmented in situ, encapsulated with barcoded beads, and processed into matched RNA and DNA methylation libraries. b,c, Per-library throughput (b) and estimated library-preparation cost per cell (c) compared with existing joint methylome–transcriptome assays. d, Enzymatic C-to-T conversion of unmethylated cytosines and retention of modified cytosines (5mC + 5hmC). e,f, Human–mouse species-mixing analysis based on DNA methylation reads (e) and RNA unique molecular identifiers (UMIs) (f). Percentages indicate human, mixed and mouse classifications. g,h, RNA (g) and DNA methylation (h) UMAP embeddings of AML01 bone marrow cells, colored by cell type. i,j, Detected genes (i) and covered CpGs (j) per AML01 cell. AML, acute myeloid leukemia; DNAm, DNA methylation; UMAP, uniform manifold approximation and projection.

Droplet-MT processed approximately 10,000 cells per library, extending joint profiling beyond the lower-throughput plate formats of scNMT-seq, snmCAT-seq and Camellia-seq^25,29–32^ (Fig. 1b; Supplementary Table 1). By pooling many cells in a single reaction, it substantially reduces reagent costs. The estimated library-preparation cost was approximately US$0.50 per cell, compared with US$20 for Camellia-seq; these estimates concern library preparation rather than total sequencing expenditure (Fig. 1c). Internal oligonucleotide controls showed 99.2% conversion of unmethylated cytosines, with only 3.8% conversion at protected modified-cytosine positions (Fig. 1d). Human-mouse mixing experiments produced predominantly species-specific profiles in both modalities, with mixed-species classifications of 3.32% for DNA and 4.35% for RNA (Fig. 1e,f). The nucleus-based workflow similarly yielded few mixed-species profiles (Extended Data Fig. 1c,d).

To assess performance in heterogeneous primary tissue, we applied the whole-cell workflow to a fresh AML bone marrow aspirate, AML01. RNA profiles resolved progenitor, monocyte, lymphoid, dendritic and plasma-cell populations, while DNA methylation provided a complementary representation of the major populations (Fig. 1g,h; Extended Data Fig. 1e). Droplet-MT also achieved medians of approximately 3,000 detected genes and one million covered CpGs per cell. Notably, its methylome coverage exceeded that of existing high-throughput methylome-only methods (Supplementary Table 1)^33,34^, providing the dense CpG sampling required for epimutation-based lineage reconstruction (Fig. 1i,j). Thus, the droplet workflow increased cellular scale while retaining the genome-wide CpG information needed for lineage-resolved analysis.

### Genome-wide epimutations resolve clonal diversity and lineage output in healthy blood

We first tested mutation-independent clonal reconstruction in healthy blood. Using the nucleus-based workflow, sorted T cells, B cells and monocytes from two donors (HB10, 17 years; F12F, 63 years) yielded 3,951 paired transcriptome–methylome profiles that passed quality control for both modalities (Fig. 2a; Extended Data Fig. 2b). Cells from both donors could be pooled within a library and demultiplexed using SNP-associated CpGs, with donor assignments supported by independently processed reference cells of known origin (Extended Data Fig. 2a,c).

**Fig. 2.**
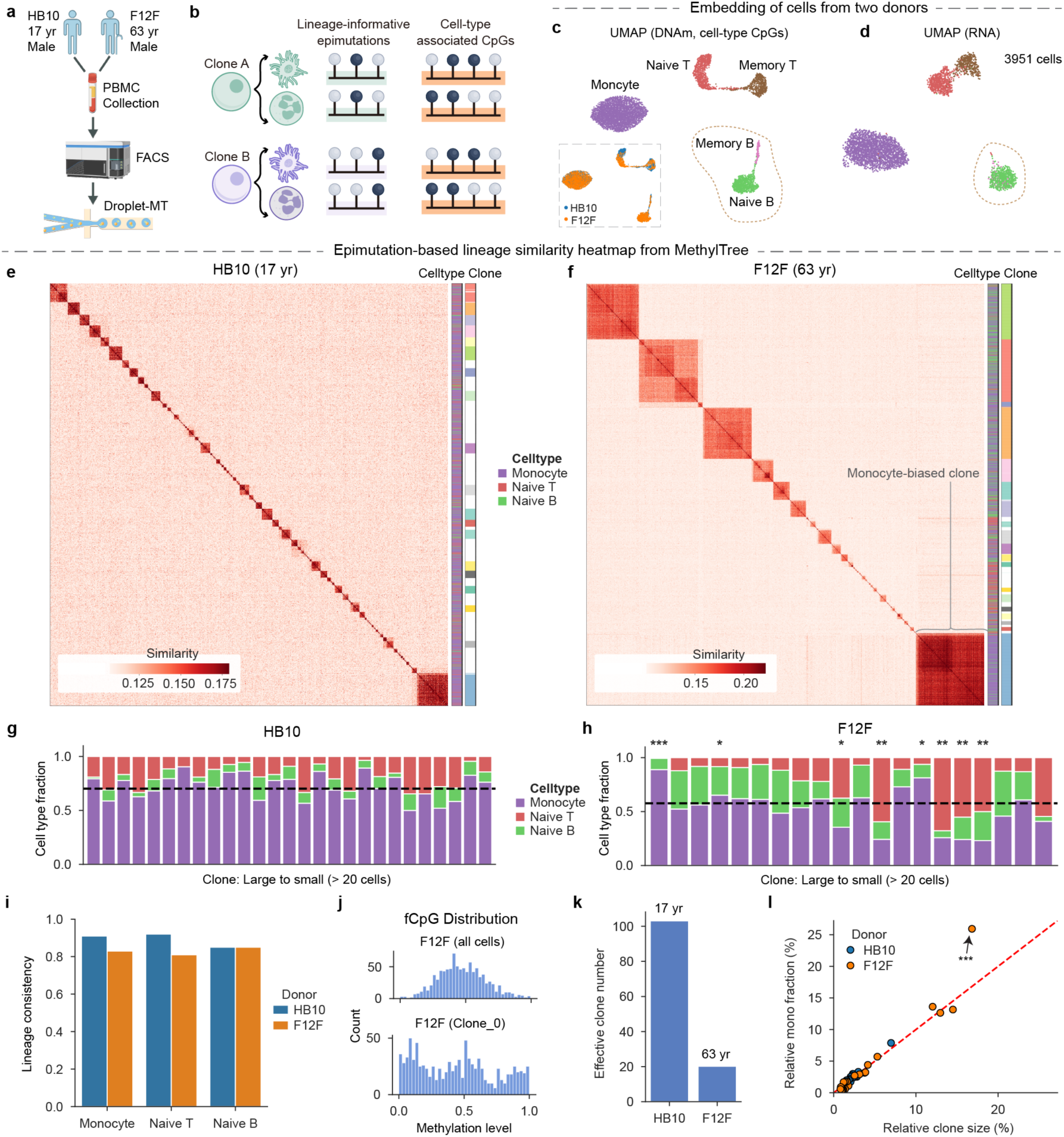
Droplet-MT reconstructs endogenous clonal architecture in healthy human blood. a, PBMC subset sorting and Droplet-MT profiling of HB10 and F12F. b, Schematic of cell-type-associated CpGs and lineage-informative epimutation CpGs. c,d, DNA methylation (c) and Harmony-integrated RNA (d) UMAP embeddings colored by cell type; inset in c shows donor identity. e,f, MethylTree cell–cell lineage similarity matrices for HB10 (e) and F12F (f), with cell-type and clone annotations. g,h, Cell-type composition of clones containing more than 20 cells in HB10 (g) and F12F (h); dashed lines show donor-level monocyte fractions. i, Concordance between clone assignments inferred from mixed cell types and those inferred within each cell type. j, fCpG methylation-state distributions in all F12F cells and in the largest F12F clone. k, Effective clone numbers of HB10 and F12F. l, Relative clone size versus the fraction of all monocytes contributed by each clone; dashed line indicates proportional monocyte output (x=y). In h and l, asterisks indicate significant deviation from donor-level cell-type fractions: *, q<0.05; **, q<0.01; ***, q<0.001 (two-sided Fisher’s exact test with Benjamini–Hochberg correction).

We analyzed the methylome through separate feature sets for cellular state and ancestry^27,28^. CpGs varying consistently between cell types were used for state embeddings, whereas CpGs with high within-cell-type but low between-cell-type variability were used for MethylTree reconstruction (Fig. 2b; Methods). The state-associated CpGs resolved the expected immune populations and separated naive from memory B cells more clearly than the RNA embedding in these nucleus-derived datasets (Fig. 2c,d; Extended Data Fig. 2g-i). CpG-based embeddings revealed shared cell-type structure across donors, whereas embeddings based on 5-kb genomic bins showed stronger donor separation (Extended Data Fig. 2d,e).

For ancestry analysis, we excluded memory B and T cells, whose global hypomethylation was consistent with activation-associated methylation remodeling^35,36^ (Extended Data Fig. 2f). MethylTree reconstruction of high-coverage cells resolved discrete similarity blocks in both donors (Fig. 2e,f; Methods). Individual blocks contained monocytes, naive T cells and naive B cells in varying proportions, linking cells across mature hematopoietic identities (Fig. 2g,h). Assignments inferred jointly across cell types were substantially concordant with those inferred within each cell type (Fig. 2i). In addition, the largest F12F clone showed a W-shaped distribution of fluctuating-CpG (fCpG) methylation states that was lost on averaging across the donor’s cells^22,23^ (Fig. 2j). These observations supported interpreting the blocks as ancestry groups, consistent with our earlier comparison of methylation-defined HSC groups with DARLIN barcodes in mice^25,28^ (Extended Data Fig. 2j).

The inferred clonal organizations differed markedly between donors. HB10 contained many small clones, whereas F12F included several expanded clones; their effective clone numbers were 103 and 20, respectively (Fig. 2e,f,k). Among clones containing more than 20 cells, eight F12F clones showed significant myeloid or lymphoid enrichment relative to the donor-specific background, whereas no HB10 clone met the same criterion after multiple-testing correction (Fig. 2g,h). The largest F12F clone accounted for 17% of analyzed cells but contributed 26% of monocytes, demonstrating disproportionate monocyte output (Benjamini-Hochberg-adjusted Fisher’s exact test, *q* = 10^-53^; Fig. 2h,l). Several F12F clones were lymphoid-enriched, indicating that clonal expansion was not uniformly associated with myeloid output. Although one donor per age group cannot establish an age effect, these analyses demonstrate that genome-wide epimutations can resolve clonal expansion and identify heterogeneous contributions from small and expanded lineages without using predefined somatic variants to distinguish them.

### High-depth clonal references enable assignment of ultra-shallow methylomes

We next separated the methylome coverage needed to discover clonal structure from that needed to track an established lineage. Pooling cells within a reference clone increases the number of covered CpGs, providing a denser comparison profile than an individual reference cell. We therefore aggregated high-depth cells into clone-specific pseudobulks and selected CpGs that varied between these references. Each query cell was compared with the reference profiles over shared CpGs and assigned to its highest-scoring clone only when the standardized similarity exceeded a specified threshold; otherwise, it remained unassigned (Fig. 3a; Methods).

**Fig. 3.**
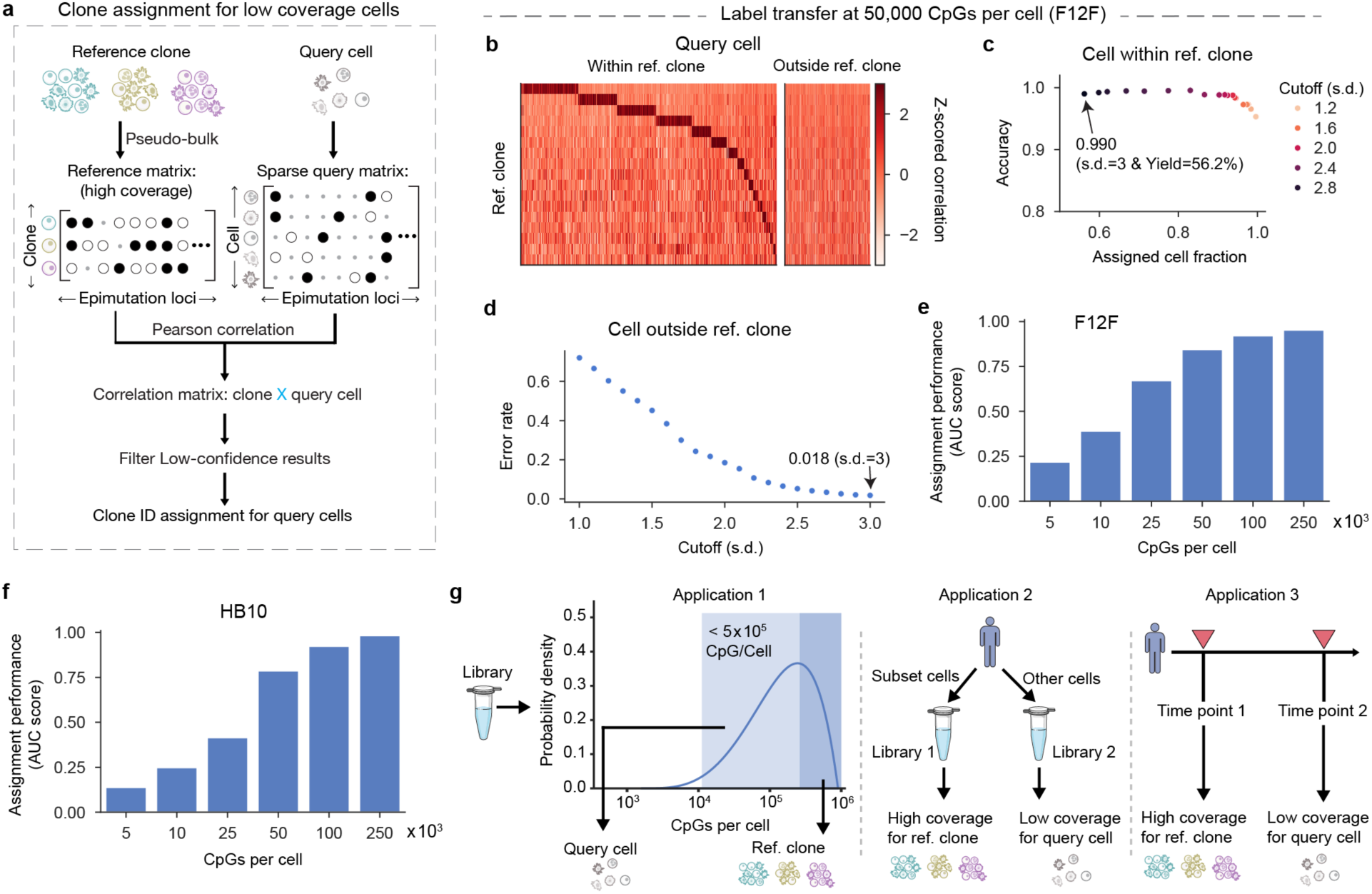
Reference-based clone-label transfer for low-coverage methylomes. a, Schematic of label transfer from high-depth reference clones to low-coverage query cells. b–d, F12F benchmark after downsampling query cells to 50,000 CpGs per cell. b, Z-scored correlations between query cells and reference clone pseudobulks, with query cells separated by whether their full-depth-defined clone is represented in the reference. c, Assignment accuracy versus assigned-cell fraction across confidence cutoffs for cells from reference-represented clones. d, False-assignment rate for cells from clones absent from the reference. Arrows mark the 3-s.d. cutoff, showing assignment accuracy and yield in c and the false-assignment rate in d. e,f, Clone-assignment performance across query-cell methylome coverage in F12F (e) and HB10 (f), summarized by area under the yield–accuracy curve across a range of cutoffs. g, Example applications of label transfer: within-library depth stratification, transfer between libraries from the same sample and longitudinal clone tracking.

We evaluated this strategy across independently generated libraries from F12F. Reference pseudobulks were constructed from clones containing more than 20 cells in one library, and cells from the second library served as queries. Full-depth methylation-based assignments supplied the benchmark labels before queries were downsampled to 50,000 covered CpGs per cell, approximately 0.17% CpG coverage. Query cells whose benchmark labels were not represented in the reference provided a separate test of inappropriate assignment to known clones (Fig. 3b).

Raising the score threshold generally improved agreement among assigned, reference-represented cells while reducing the fraction assigned (Fig. 3c). At the 3-s.d. cutoff, concordance with full-depth labels was 99.0% at 56.2% yield in F12F and 97.1% at 54.8% yield in HB10 for reference-represented cells. At the same cutoff, 1.8% of reference-absent F12F cells and 3.6% of reference-absent HB10 cells received an incorrect reference label (Fig. 3c,d; Extended Data Fig. 3a-c). Thus, high concordance for represented lineages was accompanied by a measurable trade-off between assignment yield and rejection of unrepresented lineages.

Performance improved with query coverage. The area under the yield-accuracy curve for reference-represented cells was approximately 0.84 in F12F and 0.79 in HB10 at 50,000 CpGs, and approached 1 as coverage increased to 250,000 CpGs, approximately 0.87% of genomic CpGs (Fig. 3e,f; Extended Data Fig. 3d,e). The lower performance in the small-clone-rich HB10 sample at ultra-shallow coverage was consistent with less complete coverage of its reference pseudobulks. These benchmarks support a two-stage design in which a deeply profiled subset establishes the clonal reference and additional cells are sequenced more shallowly for assignment (Fig. 3g). The same strategy can link libraries or time points, provided the lineages of interest are represented in the reference.

### Clone-level genetic annotation resolves lineage-specific changes during AML therapy

Methylation-defined ancestry can also organize genetic evidence that is too sparse to interpret one cell at a time. We tested this extension in longitudinal bone marrow samples from AML02, a 58-year-old woman with acute monocytic leukemia. Samples collected at diagnosis (T0) and on day 19 (T1), 17 days after initiation of idarubicin-cytarabine therapy (IA), yielded 5,871 paired profiles after quality control (Fig. 4a,b; Extended Data Fig. 4a). RNA and DNA methylation resolved progenitor, myeloid and lymphoid populations, with global hypomethylation in the myeloid compartment relative to naive lymphocytes (Fig. 4b-d; Extended Data Fig. 4b-f).

**Fig. 4.**
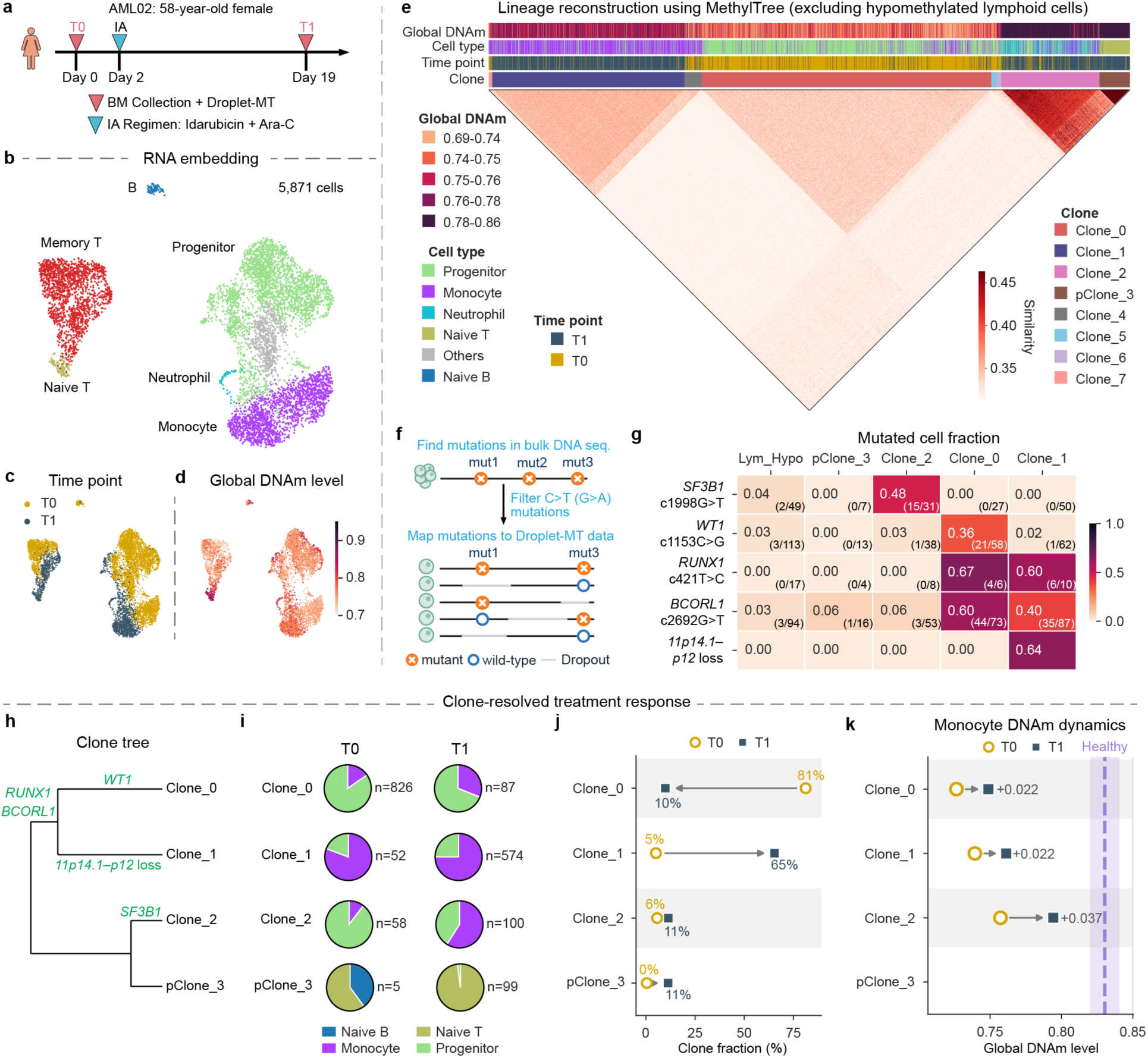
Genetic annotation links methylation-defined clones to treatment-associated changes in AML. a, AML02 sampling and idarubicin–cytarabine treatment timeline. Ara-C, Cytarabine. b–d, RNA UMAP embeddings colored by cell type (b), time point (c) or global DNA methylation level (d). e, MethylTree lineage-similarity matrix after excluding hypomethylated lymphoid cells, with annotations for DNA methylation, cell type, time point and inferred clone. pClone denotes a higher-level methylation-defined compartment containing subclonal structure. f, Strategy for querying bulk-targeted mutations in Droplet-MT DNA reads and aggregating evidence by methylation-defined clone. g, Clone-level mutation fraction among callable cells; values in parentheses indicate mutant-positive and callable cells. h, Methylation-defined clone tree annotated with clone-associated mutations and copy-number alterations. i, Cell-type composition of major clones at T0 and T1; neutrophils were excluded from this comparison because they were not sorted at T0. j, Relative clone abundance among analyzed cells at T0 and T1, without pooling-ratio adjustment. k, Global DNA methylation changes in clone-matched monocytes; labels show T1−T0 differences, and the dashed purple line indicates the healthy-monocyte mean from HB10 and F12F.

After excluding hypomethylated lymphocytes, MethylTree identified eight ancestry groups (Fig. 4e; Extended Data Fig. 4g). These included a lymphoid compartment with finer subclonal structure, denoted pClone_3; we use pClone for such higher-level, polyclonal compartments. To genetically annotate these groups despite sparse coverage at individual loci, we aggregated allele evidence across cells within each group, increasing effective locus coverage and reducing the influence of random library and sequencing errors. Candidate mutations identified by a clinical 90-gene myeloid panel were queried in existing Droplet-MT DNA reads without generating additional targeted single-cell genotyping libraries (Fig. 4f). C-to-T/G-to-A substitutions were excluded because of conversion ambiguity. For the remaining variants, we summarized the clone-level mutation fraction (CMF) as the number of mutation-positive cells divided by the number with callable evidence at that locus; cells lacking coverage did not contribute to the denominator.

The major methylation-defined branches carried distinct genetic signatures. *BCORL1* c.2692G>T and *RUNX1* c.421T>C were enriched in Clone_0 and Clone_1, with CMFs of 0.40-0.67. *WT1* c.1153C>G was enriched in Clone_0 (CMF 0.36), whereas *SF3B1* c.1998G>T marked Clone_2 (CMF 0.48; Fig. 4g). A branch-associated 11p14.1-p12 loss further distinguished Clone_1 (Extended Data Fig. 4i). These alterations provided independent genetic support for distinctions among the major branches and annotated the three principal leukemia-associated groups without requiring successful mutation detection in every member cell (Fig. 4h).

Ancestry and cellular-state annotations then separated two components of the post-treatment population shift. Within the analyzed clonal population, Clone_0 decreased from 81% to 10%, whereas Clone_1 increased from 5% to 65% (Fig. 4j). At the same time, Clone_0 and Clone_2 shifted from progenitor-dominant toward greater monocyte representation, while Clone_1 remained predominantly monocyte-like (Fig. 4i). The increased monocyte representation therefore accompanied both a change in the balance between clones and a change in state composition within existing ancestry groups. By linking these lineage-specific phenotypes and treatment-associated changes to genetic alterations, our framework enables investigation of genotype–phenotype relationships, which will require larger cohorts to establish^37–39^.

Clone-matched monocytes from the three major leukemia-associated groups also showed modest post-treatment increases in global methylation while remaining hypomethylated relative to healthy monocytes (Fig. 4k). In parallel, pClone_3, which was rare at diagnosis, became more prominent after treatment and contained multiple finer lineages composed predominantly of naive B and T cells (Fig. 4i,j; Extended Data Fig. 4j). Thus, the same reconstruction captured persistence and changing cellular output of genetically annotated AML-associated groups alongside increased representation of a polyclonal lymphoid compartment.

### Droplet-MT resolves polyclonal recovery alongside leukemia-associated persistence

We next examined whether this shared ancestry framework could resolve the organization of healthy hematopoietic recovery rather than only track a leukemia-associated genotype. AML03 was a 49-year-old man with *CBFB::MYH11*-positive AML. Standard-depth paired profiles were obtained at diagnosis (T0) and on day 39 (T1), after initiation of idarubicin, cytarabine and venetoclax (IAV) on day 2. A further sample on day 194 (T2), after continued therapy, was profiled at ultra-shallow methylome coverage (Fig. 5a). RNA profiles spanning the three time points resolved myeloid, erythroid and lymphoid states, with complementary information from DNA methylation (Fig. 5b-d; Extended Data Fig. 5a-f).

**Fig. 5.**
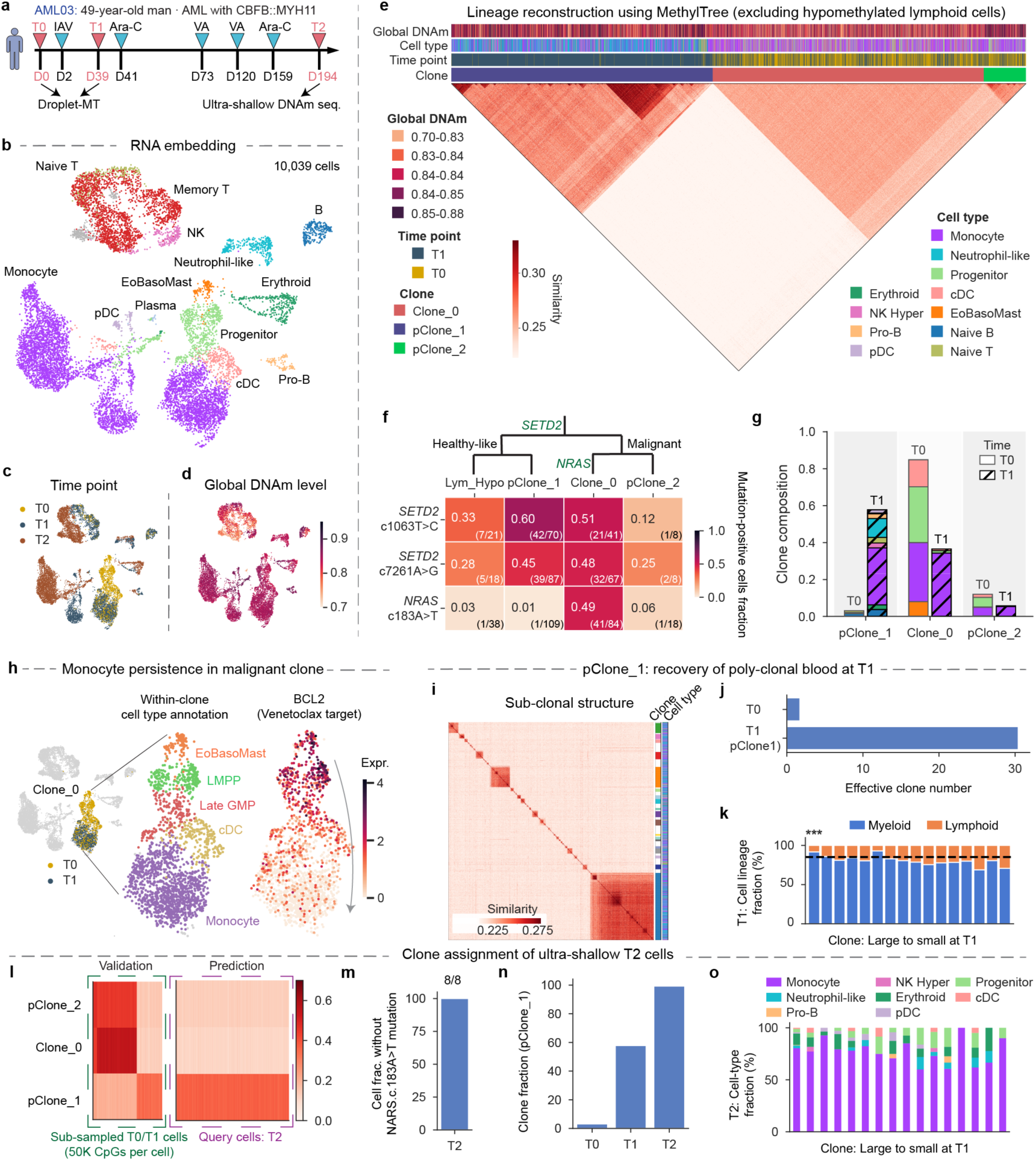
Droplet-MT distinguishes malignant persistence from polyclonal hematopoietic recovery. a, AML03 treatment and sampling timeline. Standard-depth Droplet-MT was performed at T0 and T1, and ultra-shallow DNA methylation profiling was performed at T2. b–d, RNA UMAP embeddings colored by cell type (b), time point (c) or global DNA methylation level (d). e, MethylTree lineage-similarity matrix of AML03 after excluding hypomethylated lymphoid cells; otherwise, same as Fig. 4e. f, Clone-level mutation fraction among callable cells, with a variant-annotated topology shown above. g, Relative abundance and cell-type composition of major clones at T0 and T1. h, RNA UMAPs highlighting Clone_0 cells, within-clone cell-state annotations and BCL2 expression. i, Subclonal MethylTree reconstruction of pClone_1. j, Effective clone numbers for T0 cells and T1 pClone_1 cells. k, Myeloid-vs-lymphoid lineage composition of pClone_1 subclones at T1. Myeloid cells: EoBasoMast, Progenitor, Monocyte, Neutrophil-like, cDC, pDC; lymphoid cells: NK Hyper, Naive B, Naive T, Pro-B. Asterisks indicate significant deviation from donor-level cell-type fractions (same as Fig. 2h). l, Label-transfer heatmap assigning ultra-shallow T2 cells to T0–T1 reference clones. m, Fraction of callable T2 pClone_1 cells without detected NRAS c.183A>T mutant reads. n, Relative abundance of pClone_1 at T0, T1 and T2 after excluding hypomethylated lymphoid cells. o, Predicted cell-type composition of pClone_1 subclones at T2. IAV: Idarubicin+Ara-C+Venetoclax; VA: Venetoclax+ Ara-C; NK Hyper, hyper-methylated NK cell; EoBasoMast, eosinophil/basophil/mast cell; cDC, conventional dendritic cell; pDC, plasmacytoid dendritic cell; Pro-B, progenitor B cell.

T0-T1 lineage reconstruction identified three major ancestry compartments: Clone_0, pClone_1 and pClone_2 (Fig. 5e; Extended Data Fig. 5g,h). Clone_0 formed a coherent block enriched for malignant cells at diagnosis, whereas pClone_1 contained extensive finer structure and was enriched after treatment (Fig. 5e,i). Genetic annotation distinguished a leukemia-enriched alteration from broadly distributed variants. *NRAS* c.183A>T was detected in 41 of 84 callable Clone_0 cells (CMF 0.49), compared with 1 of 109 callable pClone_1 cells (Fig. 5f). The adjacent pClone_2 had limited NRAS support (1 of 18 callable cells). By contrast, both *SETD2* variants, c.1063T>C and c.7261A>G, were distributed across the three compartments and hypomethylated lymphocytes and therefore did not specifically identify the leukemia-enriched lineage. Clone_0 and the adjacent pClone_2 were not only closely related in the epimutation-inferred lineage structure, but also exhibited similar transcriptional and epigenetic states that were distinct from pClone_1 (Extended Data Fig. 5f,i). These observations identified Clone_0 as an NRAS-enriched malignant lineage and supported interpreting pClone_2 as a putatively malignant compartment despite its limited direct mutation support.

Clone_0 and the adjacent pClone_2 decreased in relative abundance after treatment, with their residual populations predominantly occupying monocyte-like states (Fig. 5g,h). Residual Clone_0 monocytes showed low *BCL2* expression, consistent with previous observations of monocytic states in venetoclax-treated AML^40,41^ (Extended Data Fig. 5j). In contrast, pClone_1 became the dominant compartment at T1. Its effective clone number was approximately 30, substantially greater than that of the pretreatment population dominated by Clone_0 (Fig. 5i,j). Individual pClone_1 subclones contributed differently to myeloid and lymphoid lineages; notably, the largest subclone, pClone1_0, showed significant myeloid bias, resembling the clone-to-clone lineage heterogeneity observed in the older healthy donor F12F (Fig. 5k). Its high clonal diversity, multilineage output and depletion of the tracked *NRAS* variant supported interpretation of pClone_1 as recovery-associated hematopoiesis. Monocytes at T1 therefore arose both from the persisting *NRAS*-enriched lineage and from a diverse recovery-associated compartment, which could not be described by mutation status alone.

To follow this recovery at lower sequencing depth, we used the combined T0-T1 clonal reconstruction as the reference for T2 methylomes, which covered approximately 0.3% of CpGs per cell (Extended Data Fig. 5a). To accommodate this hierarchical clonal organization, assignment proceeded in stages: major compartments and the largest pClone_1 subgroup were identified by clustering reference-similarity profiles, followed by score-threshold transfer for the remaining subclones (Methods; Fig. 5l; Extended Data Fig. 5k-m). Both stages were evaluated using downsampled T1 cells with full-depth comparison labels (Fig. 5l; Extended Data Fig. 5l). T2 assignments were dominated by pClone_1, and no NRAS c.183A>T mutant reads were detected among callable T2 pClone_1 cells (0 of 8 callable cells; Fig. 5m,n). Reference-defined pClone_1 subclones continued to span multiple cell types at T2 (Fig. 5o). This analysis tracked both the changing abundance of the recovery-associated compartment and the cellular output of its constituent lineages using ultra-shallow T2 methylomes.

### Paired molecular profiles resolve lineage-associated monocyte states

Finally, we asked how clonal ancestry organized molecular variation within a shared cellular identity. Although AML blasts can differentiate into monocyte-like states, whether these cells molecularly converge toward healthy monocytes or retain a malignant identity remains unclear^19,20^. We compared CD14+ monocytes from healthy HB10 and F12F blood with CD14+ monocytes from the malignant groups in AML01-AML03 and the recovery-associated pClone_1 compartment in AML03. Diagnostic monocytes from AML01 predominantly belonged to one MethylTree-defined clone. RNA embeddings were donor-integrated with Harmony, whereas DNA methylation embeddings were generated without cross-sample integration. In both analyses, malignant monocytes remained distinct from healthy and recovery-associated groups (Fig. 6a-c). RNA profiles of malignant monocytes from different AML patients partly overlapped, whereas their methylation profiles retained stronger patient-associated structure. Within AML02, monocyte RNA profiles shifted after treatment; the corresponding AML03 leukemia-associated populations showed less separation by time point (Fig. 6b).

**Fig. 6.**
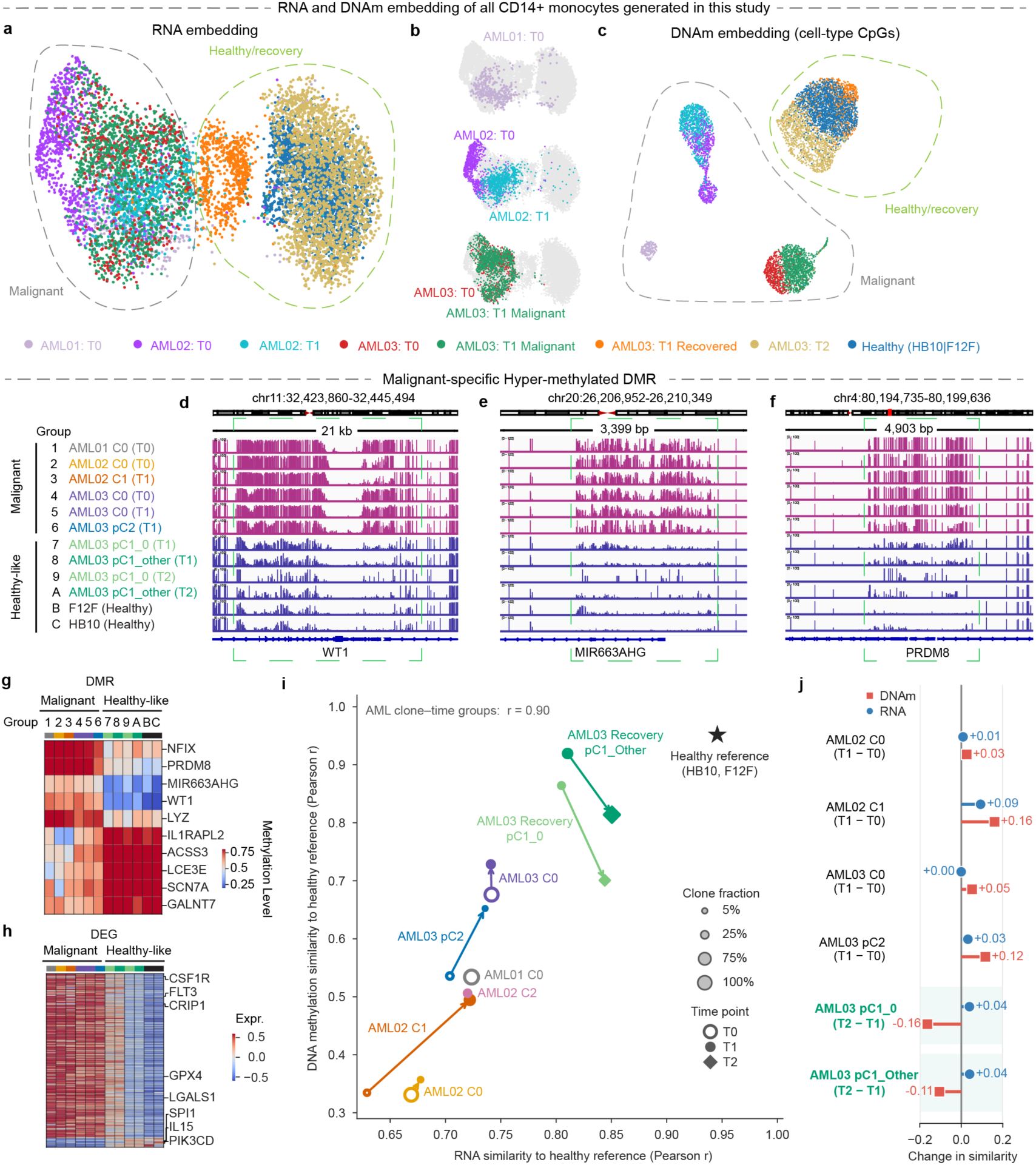
Lineage-defined CD14+ monocytes reveal malignant identity and treatment remodeling. a, Harmony-integrated RNA UMAP of CD14+ monocytes from all donors, colored by donor and sampling group. b, RNA UMAPs highlighting selected AML donor–time-point groups; other cells are gray. c, DNA methylation UMAP of the same cells based on cell-type-associated CpGs. Dashed outlines in a,c mark malignant and healthy/recovery regions. d–f, Representative malignant-specific hypermethylated differentially methylated regions (DMRs) near WT1 (d), MIR663AHG (e) and PRDM8 (f). g,h, DMR methylation levels (g) and differentially expressed genes (DEGs) (h) distinguishing malignant from normal-like CD14+ monocytes. i, RNA and DNA methylation similarity of clone–time-point groups to the healthy reference; symbol size indicates observed clone fraction without pooling-ratio adjustment, and arrows connect longitudinally related groups. The r value indicates Pearson correlation between RNA and DNA methylation similarity. j, Changes in RNA and DNA methylation similarity to the healthy reference between consecutive time points. Positive values indicate increased similarity to healthy monocytes. Recovery-associated cells are shown separately for the dominant pClone1_0 and the remaining pClone_1 subclones.

Consensus comparisons across donor-, clone- and time-point-resolved monocyte pseudobulks identified shared molecular abnormalities in malignant lineages. Recurrent differentially methylated regions (DMRs), including hypermethylated regions near *WT1*, *MIR663AHG* and *PRDM8*, were shared across donors and retained after treatment in longitudinally sampled lineages (Fig. 6d–g). Within AML03, these regions also distinguished persistent malignant monocytes from recovery-associated monocytes, providing a within-donor comparison of epigenetic states across distinct ancestry groups. Malignant monocytes also showed higher expression of *CSF1R*, *FLT3* and *CRIP1*, together with enrichment of mTORC1 signaling, unfolded protein response and reactive oxygen species pathways (Fig. 6h; Extended Data Fig. 6a). Thus, monocytes from distinct malignant lineages shared transcriptional and epigenetic abnormalities, with regional methylation differences persisting despite treatment-associated changes in cellular state.

We then quantified changes relative to healthy monocytes using these pseudobulks. For each modality, similarity was defined as the mean Pearson correlation to the separate HB10 and F12F healthy-monocyte profiles (Methods). RNA and methylation similarities were broadly concordant across AML clone-time-point groups (r = 0.90), with recovery-associated pClone_1 closest to the healthy references (Fig. 6i). However, changes between time points differed by lineage and modality. After treatment, AML02 Clone_1 showed increases of 0.09 in RNA similarity and 0.16 in methylation similarity, compared with smaller increases in Clone_0 (Fig. 6j). Treatment-associated expression and methylation features in AML02 also shifted toward the healthy-monocyte patterns (Extended Data Fig. 6b-e). AML03 Clone_0 showed little change in RNA similarity but an increase of 0.05 in methylation similarity. AML03 pClone_2 showed increases of 0.03 in RNA similarity and 0.12 in methylation similarity. Within the recovery-associated compartment, both the dominant pClone1_0 and the remaining subclones increased RNA similarity by 0.04 from T1 to T2, while methylation similarity decreased by 0.16 and 0.11, respectively (Fig. 6j). Together, these trajectories show that treatment-associated movement toward a healthy monocyte state is lineage specific and can be discordant between transcriptional and epigenetic layers, while malignant monocytes retain a molecular identity distinct from recovering and healthy hematopoiesis.

## Discussion

By integrating Droplet-MT with a comprehensive bioinformatic workflow, we establish a scalable framework for relating clonal organization to molecular state in human hematopoiesis. Building on methylation-based lineage reconstruction^28^, the advance lies in combining genome-wide CpG sampling, paired RNA profiling and depth-efficient clone tracking within a droplet workflow. Its organizing principle is that ancestry is inferred from epimutations, whereas somatic mutations provide genetic annotation when informative variants are available. Lineage analysis therefore remains possible in cells without a callable driver mutation, while retaining genetic context for malignant populations. This division of roles allows normal and leukemia-associated hematopoiesis to be examined within the same analytical framework.

Throughput and sequencing depth address different constraints on these studies. The droplet workflow supports profiling at approximately 10,000-cell library scale, recovering thousands of genes and one million CpGs per cell for both state analysis and lineage reconstruction. Across this study, we generated approximately 21,000 high-quality single-cell profiles from healthy blood and leukemia, integrating transcriptomic, epigenetic and clonal information into a resource for studying human hematopoiesis. Separating discovery from tracking then reduces the sequencing requirements for additional cells. In downsampling benchmarks, cells from reference-represented clones could be assigned at approximately 0.2% CpG coverage. Assignment yield and confidence, however, depend on reference-clone size and coverage. This is particularly important in normal tissues, where small clones may be poorly represented, and during recovery, when previously rare lineages can become abundant. References may therefore require additional deep sampling rather than remaining fixed at diagnosis.

Genetic annotation reuses sequence information already present in methylome libraries. Variants identified by bulk DNA sequencing can be queried retrospectively, with allele evidence aggregated across cells within each epimutation-defined clone to increase effective locus coverage and reduce the influence of random PCR and sequencing errors. Genetic targets need not be specified before data collection, and covered coding and noncoding loci can be interrogated independently of transcript abundance or position. C-to-T/G-to-A substitutions were excluded because of conversion ambiguity; TAPS, which leaves unmodified cytosines unchanged, may alleviate this issue^42^.

The healthy-blood analyses illustrate the main opportunity offered by this approach. Small and expanded methylation-defined clones linked monocytes with naive B and T cells and differed in their contributions to these populations. Such measurements make it possible to ask whether changes in blood-cell composition are broadly shared across lineages or concentrated in a subset of clones. In particular, the older donor showed reduced effective clonal diversity and greater heterogeneity in lineage output than the younger donor, with individual clones displaying either myeloid-or lymphoid-biased contributions rather than a uniform shift toward myeloid output. However, one individual per age group cannot establish an age effect. Larger longitudinal cohorts will be required to connect these clone-resolved measurements to population-level patterns of hematopoietic aging.

The AML analyses provided a complementary application because malignant and recovery-associated populations could be examined together. In AML02, treatment was associated with changes in the relative abundance of malignant clones and in their cellular output. In AML03, leukemia-associated monocytes coexisted with a clonally diverse, multilineage compartment that became dominant after treatment. Branch-associated somatic alterations supported the genetic interpretation of malignant lineages, whereas epimutations resolved the organization of recovery-associated cells not distinguished by those mutations. The resulting information is not merely a malignant-versus-normal classification: it describes recovery as a distribution of clonal contributions. Paired molecular readouts extend the analysis beyond clonal abundance. Within lineage-defined CD14+ monocytes, malignant populations remained distinct from healthy and recovery-associated groups in both RNA and methylation profiles, and RNA and methylation states did not always remodel in parallel over treatment, highlighting the need to jointly measure both modalities in the same cell. These cases illustrate a shared framework for studying malignant persistence and hematopoietic reconstitution, but their heterogeneous treatments and sampling intervals preclude general clinical conclusions.

Beyond hematopoiesis, methylome-based lineage tracing has been demonstrated in HEK293T cells, human embryonic stem cells and human colorectal cancer^28^, supporting its broader applicability. Droplet-MT could extend such studies by linking clonal organization to transcriptional and epigenetic states during tissue maintenance, aging and regeneration. The retrospective genetic annotation of the epimutation-defined clones can be particularly useful to explore the genotype-phenotype relationship in cancer. Building on advances in spatially resolved barcode-based lineage tracing^43–45^, integration with spatial transcriptome–methylome profiling could further place human clonal relationships within their tissue context^46,47^. Together, these developments offer a path toward understanding how clonal history, molecular state and spatial organization shape the maintenance and remodeling of human tissues.

**Extended Data Fig. 1.**
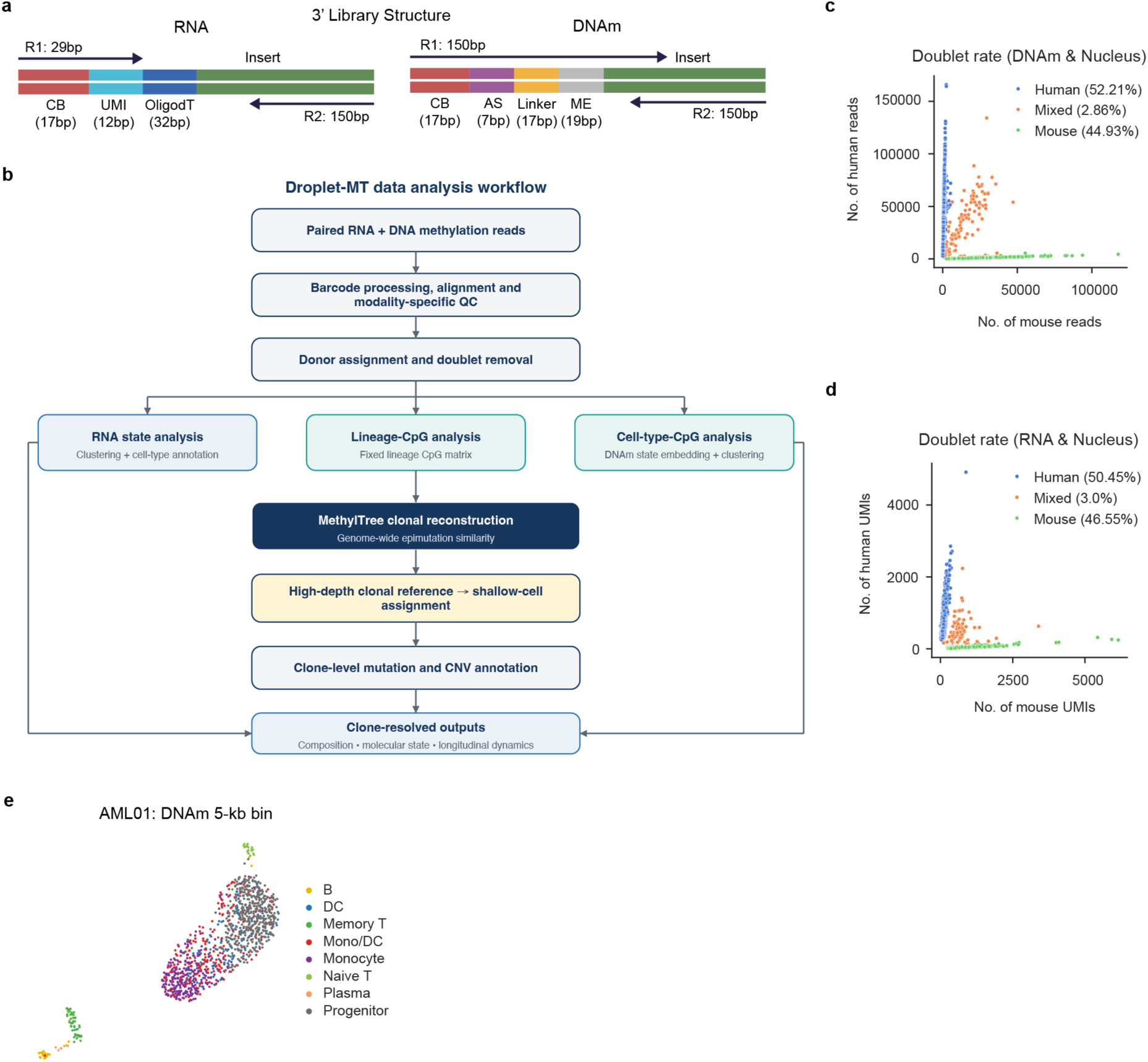
Droplet-MT library design, computational workflow and species-mixing controls. a, RNA and DNA methylation library structures generated by Droplet-MT. b, Overview of Droplet-MT preprocessing and downstream analysis. c,d, Human–mouse mixing analysis using the nucleus-based Droplet-MT workflow; otherwise, same as Fig. 1e,f. e, AML01 DNA methylation UMAP based on 5-kb genomic bins, colored by annotated cell type.

**Extended Data Fig. 2.**
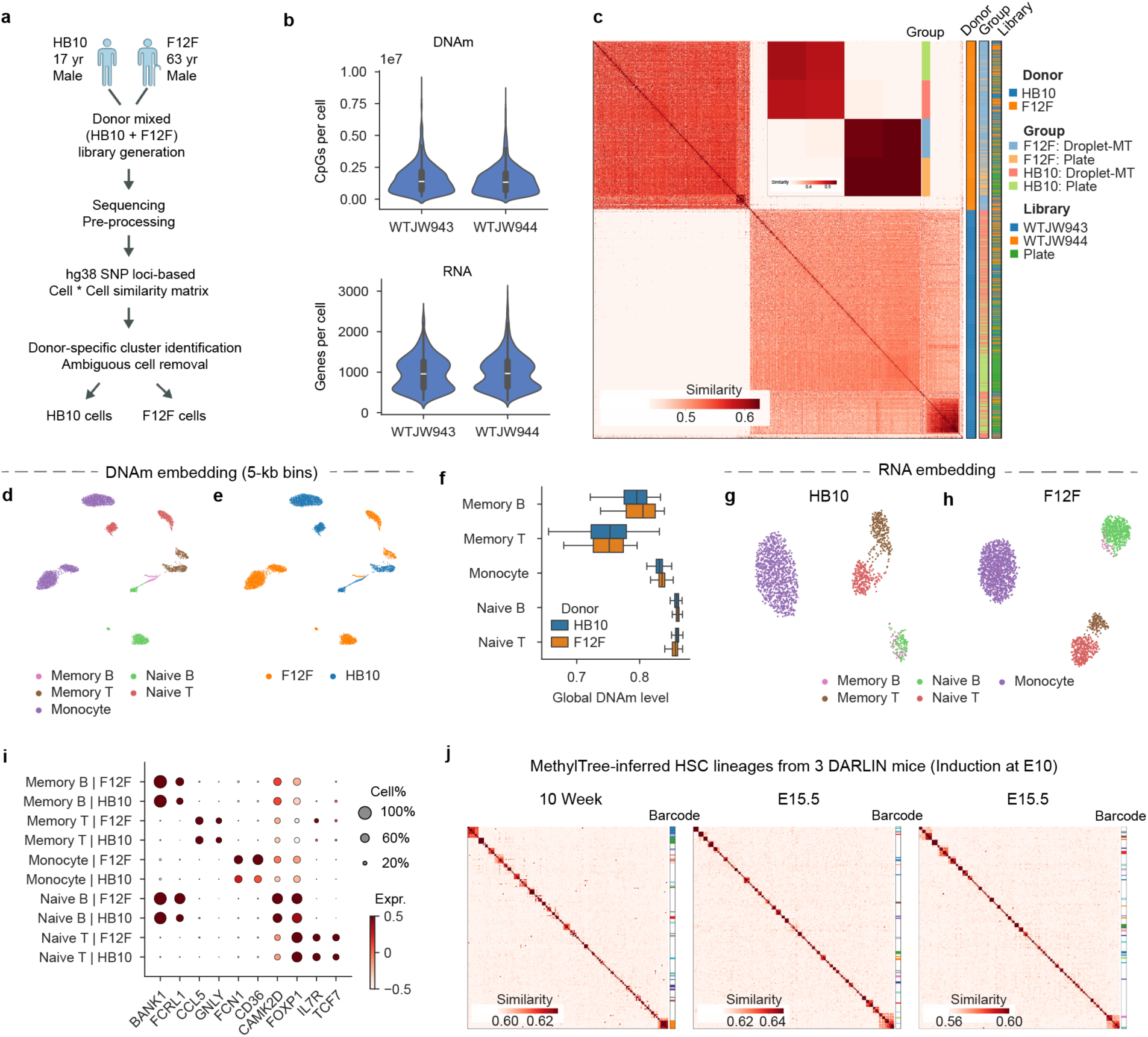
Donor demultiplexing, immune-cell states and supporting lineage analyses. a, Mixed-donor library generation and SNP-based demultiplexing of HB10 and F12F. b, Covered CpGs and detected genes per cell in two mixed-donor libraries. c, Donor-assignment validation using independent plate-based data; inset shows pseudobulk similarities. d,e, DNA methylation UMAPs based on 5-kb genomic bins, colored by cell type (d) or donor (e). f, Global DNA methylation levels across cell types. Boxes show interquartile range and median; whiskers extend to 1.5× the interquartile range, and outliers are omitted. g,h, RNA UMAP embeddings for HB10 (g) and F12F (h). i, Canonical marker-gene expression across annotated cell types. Dot size indicates the percentage of expressing cells, and color indicates scaled mean expression. j, MethylTree lineage-similarity matrices of hematopoietic stem cells (HSCs) from three independent DARLIN mice. Barcode labeling was induced at embryonic day 10 (E10), and HSCs were profiled at 10 weeks of age (left) or E15.5 (middle and right). Similarities were generated using the genome-wide CpG set (∼20 million sites). Diagonal blocks reveal methylation-defined ancestry groups of varying sizes in all three mice. Adjacent annotation bars indicate DARLIN barcode-defined clones, providing an independent comparison with the methylation-inferred structure. White indicates missing barcode assignments or small barcode-defined clones not individually colored.

**Extended Data Fig. 3.**
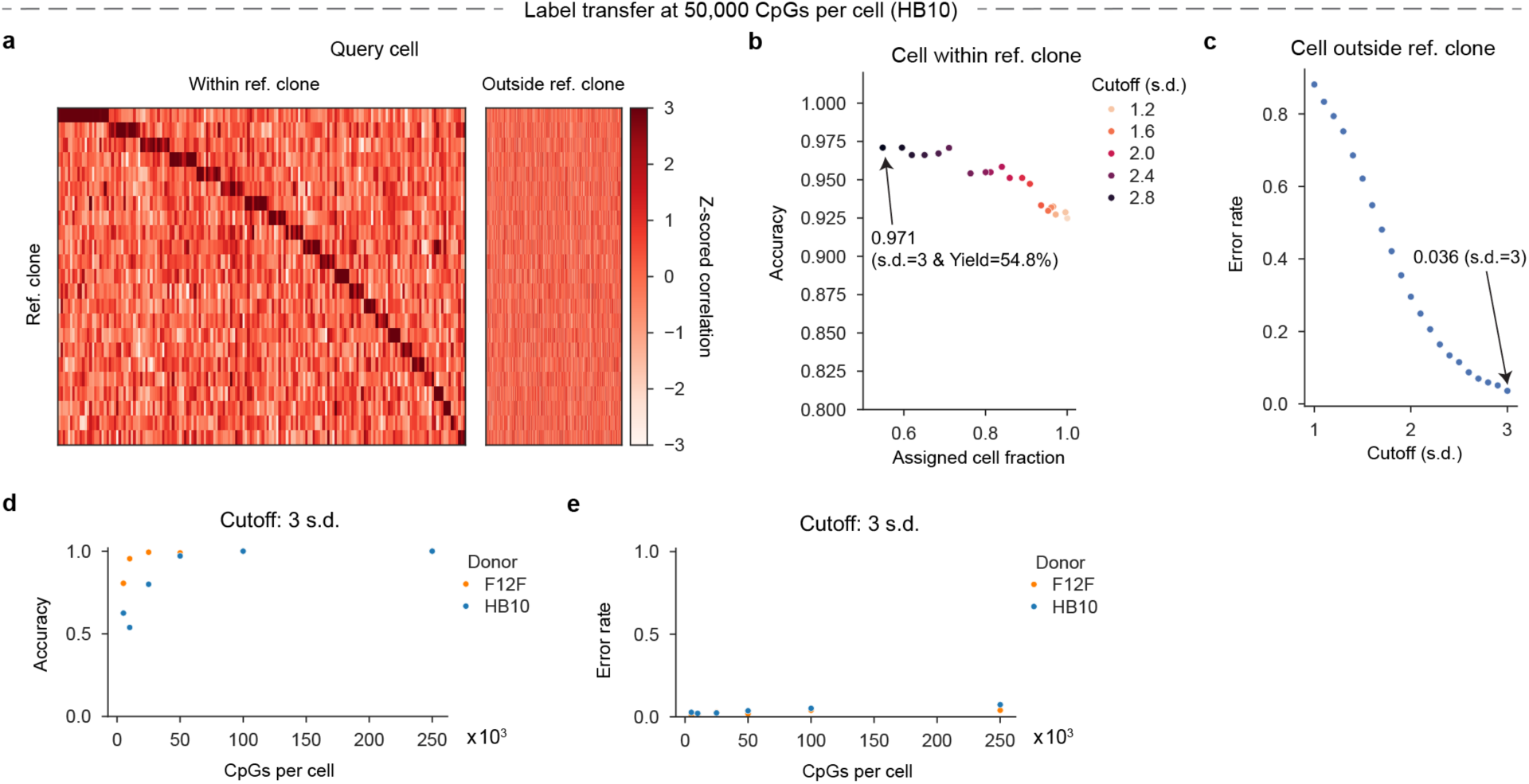
**Clone-assignment performance across query coverage and score thresholds.**a–c, HB10 benchmark after downsampling query cells to 50,000 CpGs per cell. a, Z-scored correlations between query cells and reference clone pseudobulks, separated by whether the full-depth-defined clone is represented in the reference. b, Assignment accuracy versus assigned-cell fraction across confidence cutoffs for reference-represented cells. c, False-assignment rate for cells from reference-absent clones. Arrows in b,c indicate the downstream confidence cutoff (3 s.d.). d,e, Assignment accuracy for reference-represented cells (d) and false-assignment rate for reference-absent cells (e) across query-cell coverage in HB10 and F12F using the 3-s.d. cutoff.

**Extended Data Fig. 4.**
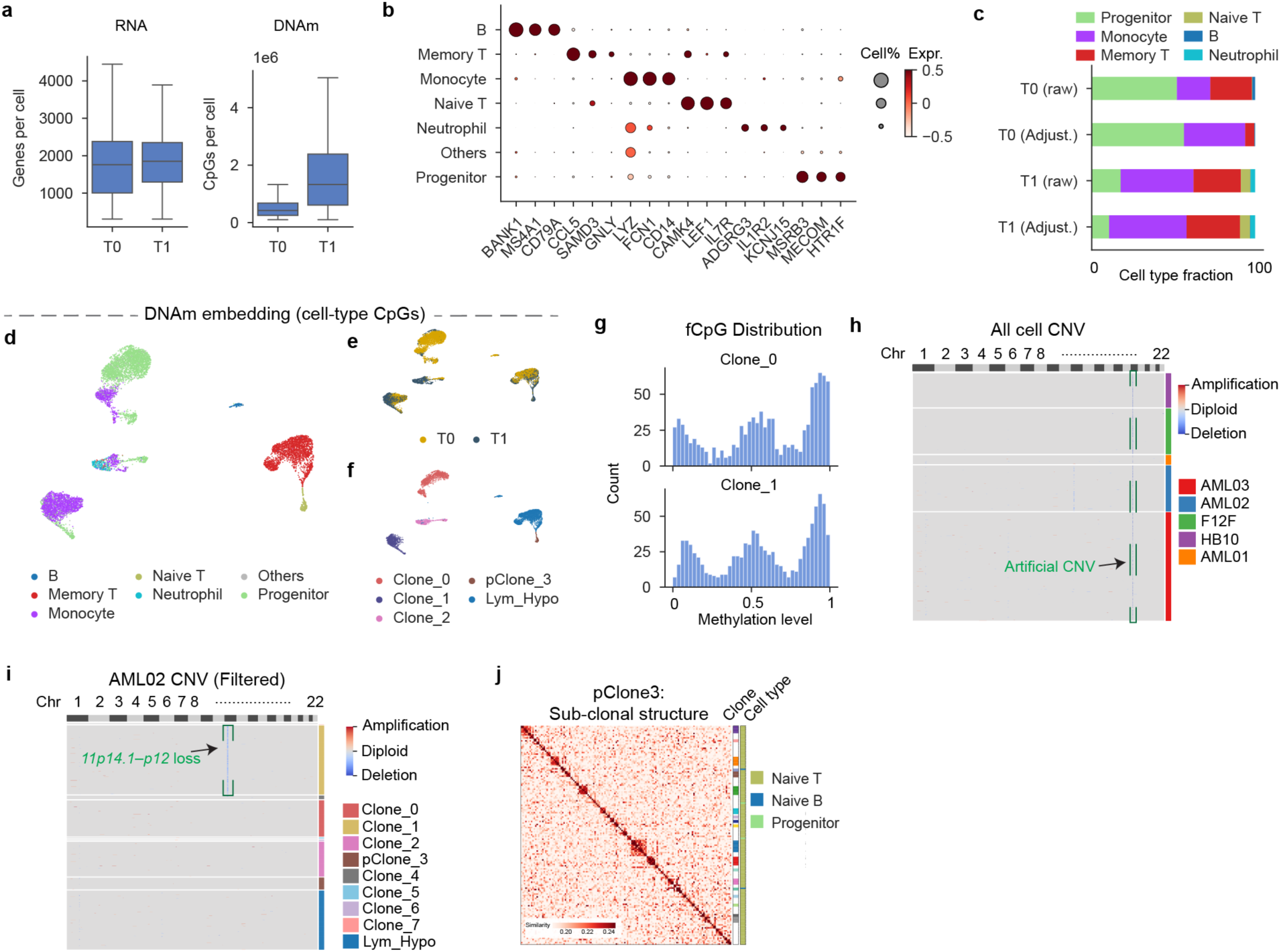
Molecular and genetic characterization of AML02 ancestry groups. a, Detected genes and covered CpGs per AML02 cell at T0 and T1. Boxes show interquartile range and median; whiskers extend to 1.5× the interquartile range, and outliers are omitted. b, Canonical marker-gene expression across annotated cell types. c, Observed and pooling-ratio-adjusted cell-type compositions at T0 and T1. Observed fractions reflect the experimentally pooled populations; adjustment using FACS proportions illustrates correction for pooling. Main-text analyses use the observed, unadjusted fractions. d–f, DNA methylation UMAPs based on cell-type-associated CpGs, colored by cell type (d), time point (e) or inferred cell group/clone (f). g, fCpG methylation-state distributions in Clone_0 and Clone_1. h, CNVpytor-predicted clone-resolved copy-number profile for healthy and AML donors, with an artificial CNV signal that showed up across all cells. i, Clone-resolved copy-number profiles in AML02, after removing the putative artifacts in h, with the 11p14.1–p12 loss indicated. j, Subclonal MethylTree reconstruction of pClone_3 with inferred-subclone and cell-type annotations. High noise here is likely due to both a smaller number of cells and a high clonal diversity.

**Extended Data Fig. 5.**
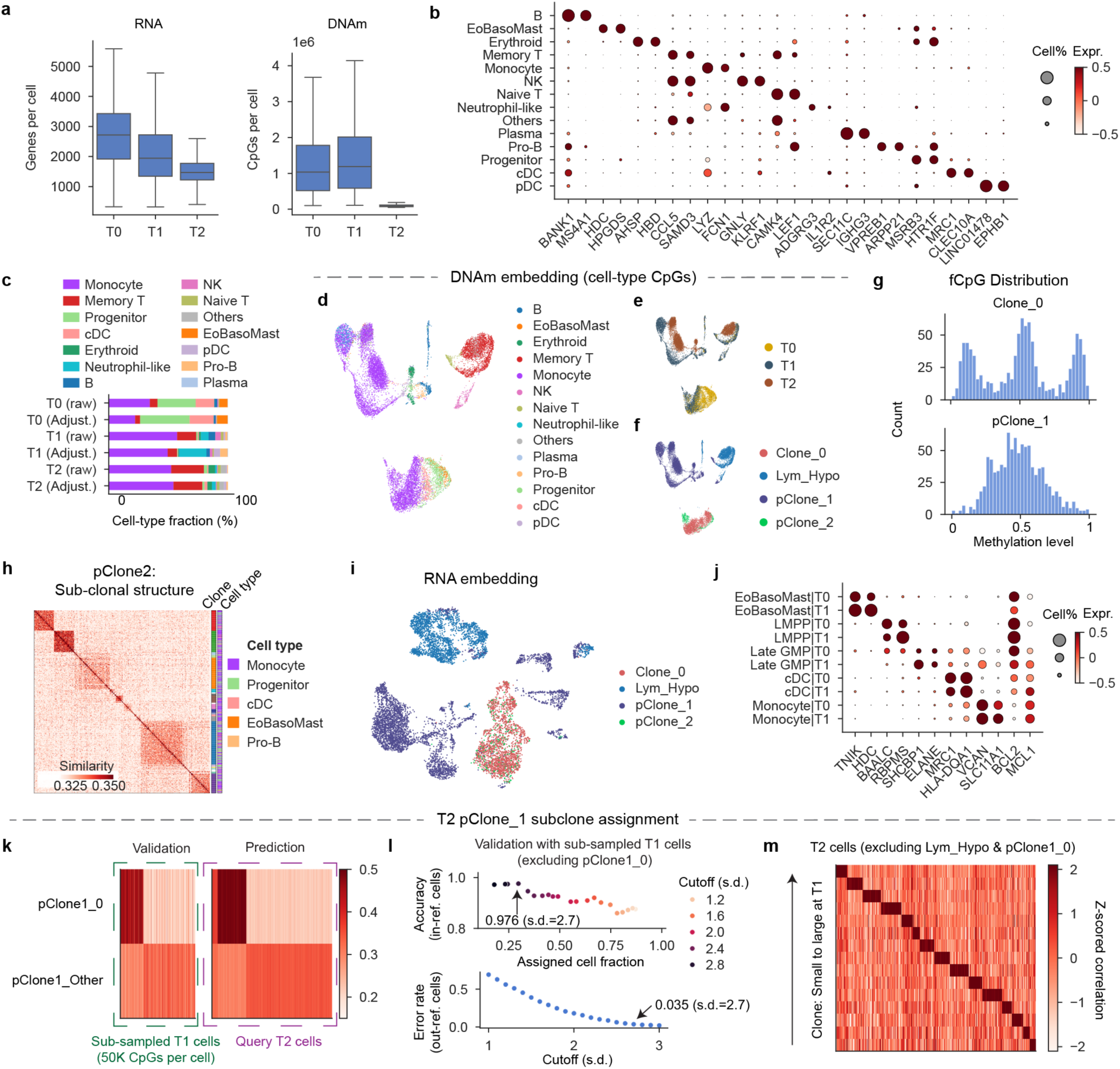
AML03 clonal structure and staged assignment of ultra-shallow methylomes. a, Detected genes and covered CpGs per AML03 cell at T0, T1 and T2. b, Canonical marker-gene expression across annotated cell types. c, Observed and FACS-pooling-ratio-adjusted cell-type compositions, shown to illustrate potential correction for experimental pooling; main-text analyses use unadjusted fractions. d–f, DNA methylation UMAPs based on cell-type-associated CpGs, colored by cell type (d), time point (e) or inferred clone/group (f). g, fCpG methylation-state distributions in Clone_0 and pClone_1. h, Subclonal MethylTree reconstruction of pClone_2. i, RNA UMAP colored by inferred clone/group. j, Marker-gene expression across Clone_0 cell states at T0 and T1. k, Assignment of ultra-shallow T2 pClone_1 cells to the dominant subclone pClone1_0, validated using downsampled T1 cells excluded from the reference pseudobulks. l, Assignment-performance validation for the remaining pClone_1 subclones using downsampled T1 cells excluded from the reference pseudobulks. m, Assignment of the remaining pClone_1 subclone labels to T2 cells after excluding hypomethylated memory lymphocytes and cells assigned to pClone1_0. Only predictions above the selected cutoff (s.d.=2.7) were shown in the heatmap.

**Extended Data Fig. 6.**
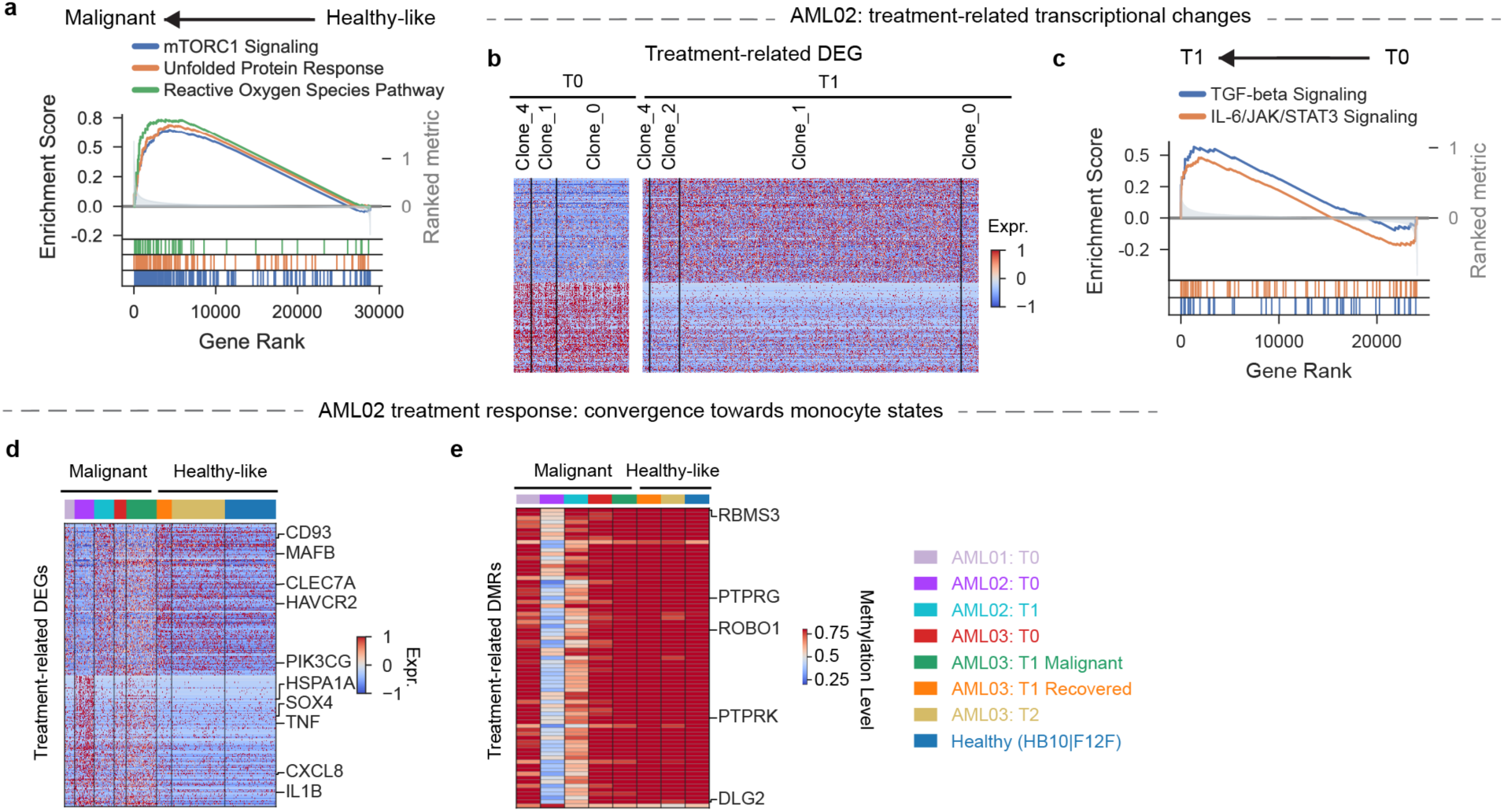
AML-associated monocyte programs and treatment-related molecular changes in AML02. a, Gene set enrichment analysis (GSEA) of pathways enriched in malignant versus healthy-like CD14+ monocytes. b, Heatmap of treatment-related DEGs across AML02 clones at T0 and T1. c, GSEA of pathways enriched at T1 relative to T0 in AML02. d,e, Heatmaps of AML02 treatment-related DEGs (d) and DMRs (e) across malignant and healthy-like CD14+ monocyte groups.

## Methods

### Human participants and ethics approval of this study

Human peripheral-blood from two healthy donors (HB10 & F12F) and bone-marrow specimens from three AML patients (AML01-AML03) were obtained with informed consent under protocols approved by Qilu Hospital of Shandong University [KYLL-2026-05-058-1]. Samples were handled in accordance with the Declaration of Helsinki and applicable institutional regulations. Full donor and library information can be found in Supplementary Table 2.

### Flow cytometry and cell sorting

Human peripheral-blood leukocytes and AML bone-marrow cells were washed and resuspended in EasySep buffer (calcium-and magnesium-free HBSS supplemented with 25 mM HEPES, 2 mM EDTA and 2% FBS). Fc receptors were blocked with Human TruStain FcX (BioLegend, 422302), followed by antibody staining. Peripheral-blood samples were stained for CD45, CD14, CD16, CD19, CD27, CD3, CD45RA and CD45RO, whereas AML samples were stained for CD14, CD34, CD45 and CD117. Cells were stained with DAPI immediately before sorting on a Sony MA900 cell sorter in Purity mode. Standard compensation and gating procedures were used to exclude debris, doublets and non-viable cells.

Peripheral-blood populations were defined as monocytes (CD45^+^CD14^+^CD16^−^), granulocytes (CD45^+^CD14^−^CD16^+^), Naive T cells (CD45^+^CD19^−^CD3^+^CD45RA^+^CD45RO^−^) and Naive B cells (CD45^+^CD3^−^CD19^+^CD27^−^). In AML samples, progenitors were defined as CD45^+^ cells expressing CD34 and/or CD117, thereby including CD34+CD117+, CD34+CD117− and CD34−CD117+ cells. For AML01, T cells, B cells, monocytes and an abnormal granulocytic population were separately collected using CD3, CD19, CD14 and CD56-based gating. For AML02, AML03 and their post-treatment samples, lymphocytes were collected from the CD45^+^ lymphocyte-scatter gate, whereas monocytes and granulocytes outside this gate were defined as CD45^+^CD34^−^CD117^−^CD14^+^ and CD45^+^CD34^−^CD117^−^CD14^−^, respectively. Sorted cells were pooled together with a sample-specific ratio to balance different populations before Droplet-MT sequencing (Supplementary Table 3). The observed cell type proportion can be corrected with this FACS-pooling ratio to give more realistic proportion (Extended Data Fig. 4c,5c). However, the plots in the main figures show the observed proportion in the single-cell data.

### Whole-cell 3′ Droplet-MT workflow

#### Whole-cell fixation and chromatin relaxation

Approximately 2 × 10^5^ sorted cells were pelleted at 500 × g for 7 min at 4 °C and resuspended in 80 µL PBS-RI (PBS containing 1 U µL^−1^ RNase inhibitor). Ice-cold methanol (400 µL) was added dropwise with gentle mixing, resulting in an approximately 80% (v/v) methanol solution. Cells were incubated for 30 min at −20 °C, during which methanol fixation simultaneously permeabilized the cells and relaxed chromatin, thereby increasing genomic DNA accessibility while preserving cellular integrity and RNA retention. Fixed cells were pelleted at 500 × g for 10 min at 4 °C, washed once in Wash Buffer A (3× SSC, 1 U µL^−1^ RNase inhibitor and 0.1% Tween-20) and once in Wash Buffer B (PBS, 1 U µL^−1^ RNase inhibitor and 0.1% Tween-20), and resuspended in PBS-RI. The resulting fixed and chromatin-relaxed cells were subsequently subjected to Tn5 tagmentation.

#### Tn5 assembly and in situ DNA tagmentation

A custom asymmetric Tn5 transposome (Tn5 Enzyme A) was assembled using three oligonucleotides:

Tn5-A: 5′-[Phos]CTGTCTCTTATACACATCT-3′

Tn5-B: 5′-[Phos]CGTCCGTCGTTGCTCGAGATGTGTATAAGAGACAG-3′

Tn5-C: 5′-GT[5mC]T[5mC]GTGGG[5mC]T[5mC]GGAGATGTGTATAAGAGA[5mC]AG-3′,

where [Phos] denotes a 5′ phosphate modification and [5mC] denotes 5-methylcytosine. Tn5-A serves as the common complementary mosaic-end oligonucleotide and was annealed to Tn5-B and Tn5-C to generate the asymmetric adapters used for Tn5 loading. The Tn5-B adapter provides the phosphorylated end used for subsequent ligation of the droplet-derived cell barcode, whereas methylated cytosines in Tn5-C preserve the adapter sequence during downstream C-to-T conversion. These fixed adapter sequences with predefined cytosine methylated states were subsequently used as internal controls for enzymatic conversion and 5mC retention.

Approximately 3 × 10^4^ fixed cells were incubated with 11 U preassembled Tn5 Enzyme A in a 60 µL reaction for 60 min at 37 °C. Tagmentation was terminated by two washes with PBS containing 0.1% Tween-20. Cells were resuspended in 50 µL PBS-RI, counted by acridine-orange/propidium-iodide staining and used immediately for droplet encapsulation.

#### Droplet barcoding and paired molecular capture

Gel beads carried separate DNA- and RNA-capture oligonucleotides encoding matched bead identities, enabling DNA and RNA molecules captured within the same droplet to be linked computationally. The paired DNA and RNA identifiers corresponded to the same bead identity but were not necessarily identical nucleotide sequences. The DNA-capture oligonucleotide contained a biotin moiety for subsequent enrichment, a 17-nt C-free cell barcode designed to remain unchanged during C-to-T conversion, and a terminal sequence for bridge-mediated ligation to tagmented genomic DNA. The RNA-capture oligonucleotide contained the corresponding 17-nt cell barcode, a 12-nt unique molecular identifier (UMI), and a poly(dT)_30_ VN sequence for capture of polyadenylated RNA. The full bead-oligonucleotide structures were (Extended Data Fig. 1a):

DNA: 5′-A[Biotin-dT]A[5mC]GA[5mC]G[5mC]T[5mC]TT[5mC][5mC]GAT[5mC]T[J]_17_TTGCTGT-3′

RNA: 5′-A[5mC]GA[5mC]G[5mC]T[5mC]TT[5mC][5mC]GAT[5mC]T[N]_17_[N]_12_(T)_30_VN-3′

where [J]_17_ denotes the 17-nt C-free DNA cell barcode, and [N]_17_ and [N]_12_ denote the RNA cell barcode and UMI, respectively. The flanking fixed sequences in the DNA-related bead oligo provided predefined cytosine positions for internal conversion-quality control.

Tagmented cells were resuspended in an 80-µL ligation and reverse-transcription mixture containing 8 µL 10× Ligation Buffer (Enzymatics, B6030L), RNase inhibitor (1 U µL^-1^), 1 mM each dATP, dTTP, dGTP and 5-methyl-dCTP (NEB, N0356S), 15 mM DTT, 3.2 µL T4 DNA ligase (Enzymatics, L6030), 5.2 µL reverse transcriptase (SeekGene, R0003802), 20 µM template-switching oligonucleotide and 0.3 µM bridge oligonucleotide (5′- AGCAACGACGGACGACAGCAA-3′). The bridge oligonucleotide hybridized the bead-derived DNA-capture sequence to the Tn5-B adapter, thereby positioning the two molecules for ligation and transfer of the bead-specific cell barcode to genomic DNA.

Cells and barcoded gel beads were co-encapsulated in droplets using a SeekOne Digital Droplet instrument (SeekGene, M001A). Within each droplet, genomic DNA fragments were ligated to bead-derived DNA barcodes while polyadenylated RNA underwent reverse transcription with the matched RNA barcode. Emulsions were incubated for 30 min at 25 °C to support DNA ligation, followed by 60 min at 42 °C for reverse transcription. Reactions were terminated with 3 µL 10% SDS, and the emulsion was broken to release the barcoded nucleic acids.

#### Separation of DNA and cDNA

Biotinylated genomic DNA was captured with M-280 Streptavidin magnetic beads (Thermo Fisher Scientific, 11205D), while the barcoded cDNA remained in the supernatant. The bead-bound DNA fraction was retained for methylome library construction, and the cDNA-containing supernatant was purified with VAHTS DNA Clean Beads (Vazyme, N411-03) and processed independently. This physical separation allowed the DNA and RNA fractions to undergo modality-specific library preparation while preserving their matched cell identities through the paired bead barcodes.

#### DNA methylation library construction

Streptavidin-bound DNA was subjected to gap filling for 20 min at 65 °C in a 50-µL reaction containing 5-methyl-dCTP in place of dCTP, thereby protecting newly synthesized adapter and barcode sequences from subsequent C-to-T conversion. DNA methylation conversion was then performed using the NEBNext Enzymatic Methyl-seq kit (NEB, E7120S), including TET2 oxidation followed by APOBEC-mediated deamination. Converted DNA was amplified for 7-9 cycles using primers 5′-CTACACGACGCTCTTCCGATCT-3′ and 5′-GTCTCGTGGGCTCG*G*A*G-3′, where asterisks denote phosphorothioate bonds, and purified at a 0.8× bead ratio. Approximately 100 ng of amplified DNA was then tagmented with Nextera XT, dual-indexed using TruSeq P5 and Nextera P7 primers for eight cycles, and purified at a 0.7× bead ratio to generate the final methylome library.

#### 3′ Transcriptome library construction

Purified cDNA was amplified for seven cycles using primers 5′-CTACACGACGCTCTTCCGATCT-3′ and 5′-AAGCAGTGGTATCAACGCAGAGT-3′, followed by purification at a 0.8× bead ratio. An aliquot of the amplified cDNA was subjected to an additional round of amplification with the same primer pair and purified at a 0.6× bead ratio. The resulting cDNA was then tagmented, dual-indexed, and purified at a 0.8× bead ratio to generate the final 3′ transcriptome library.

#### Sample-specific input configurations

Unless otherwise stated, samples were processed using the whole-cell 3′ Droplet-MT workflow described above. HB10 and F12F were instead processed from isolated nuclei using the 3′ Droplet-MT workflow. Briefly, intact cells were lysed in ice-cold nuclei isolation buffer (100 mM Tris-HCl, pH 8.0, 50 mM KCl, 10 mM DTT, 1% IGEPAL CA-630, and 1 U µL⁻¹ RNase inhibitor), and nuclei were collected by centrifugation at 500 × g for 5 min at 4 °C. Isolated nuclei were washed and subsequently subjected to the same fixation, in situ tagmentation, and droplet-barcoding procedures used for the whole-cell workflow, with RNA capture restricted to nuclear polyadenylated transcripts. AML03 T2 was processed using the SeekOne DD Single-Cell DNA Methylation + RNA Kit (SeekGene, K00901), which uses 5′ RNA capture; the DNA methylome workflow was otherwise unchanged. The workflow used for each sample is summarized in Supplementary Table 2.

#### Library quality control and sequencing

Final libraries were quantified by Qubit fluorometry, and fragment-size distributions were assessed using an Agilent 4200 TapeStation. Libraries were required to have a concentration of at least 5 ng µL^-1^, a predominant fragment-size distribution of 300-1,000 bp for transcriptome libraries and 350-1,500 bp for methylome libraries, minimal short-fragment contamination in transcriptome libraries, and minimal high-molecular-weight contamination in methylome libraries. Libraries failing these criteria were not sequenced. Both transcriptome and DNA methylome libraries were sequenced on GeneMind SURFseq Q platform, using paired-end 150-cycle sequencing. Dual 8-bp sample-index reads (i7 and i5) were collected for both transcriptome and methylome libraries to enable library demultiplexing.

#### Evaluation of cytosine-conversion efficiency

Fixed sequences introduced by the Tn5 and bead oligonucleotides were used as internal controls for enzymatic cytosine conversion. Predefined unmethylated cytosines in the AS, Linker and ME regions were expected to convert to thymine, whereas corresponding 5mC-protected positions were expected to remain as cytosine (Extended Data Fig. 1a). Conversion efficiency and 5mC retention were calculated from these diagnostic positions using SeekSoulMethyl v2.1.2.

#### Species-mixing experiment for doublet evaluation

Granulocytes, monocytes, and lymphocytes were isolated separately from human peripheral blood and C57BL/6J mouse blood, yielding ∼4 × 10^5^ cells per species. Human and mouse cells were fixed and tagmented separately, mixed immediately before droplet encapsulation, and then processed together for library construction.

These reads were aligned to a combined hg19-mm10 reference with Bismark (v0.24.0). For each cell barcode, species fractions were calculated from uniquely mapped DNA methylation reads and RNA UMI counts. Barcodes were classified as human if >90% of DNA methylation reads or >75% of RNA UMIs mapped to hg19, and as mouse if >90% of DNA methylation reads or >90% of RNA UMIs mapped to mm10. Thresholds were chosen based on the observed species-assignment distributions. Barcodes not meeting either species criterion were classified as mixed-species and considered putative interspecies doublets.

#### Overview of Droplet-MT data analysis

Droplet-MT data were processed using an integrated analysis workflow that leveraged paired transcriptomic and DNA methylation measurements from the same cells (Extended Data Fig. 1b). RNA and DNA methylation reads were first processed independently using modality-specific pipelines for barcode extraction, read alignment and quality control. For donor-multiplexed libraries, cells were assigned to individual donors using SNP-associated CpG sites, and putative doublets were identified and removed using complementary information from the RNA and DNA methylation modalities.

Following preprocessing, the paired data were analyzed along three complementary axes. RNA profiles were used for transcriptional dimensionality reduction, clustering, cell-type annotation and characterization of molecular states. DNA methylation profiles were analyzed with MethylTree (v.0.2.4) using two distinct sets of CpG features designed to capture cell-state-and lineage-associated variation. Cell-type-associated CpGs were used for methylation-based dimensionality reduction, clustering and characterization of epigenetic cell states, whereas lineage-informative CpGs were used by MethylTree to quantify epimutation similarity and reconstruct clonal relationships. High-coverage cells with robust de novo clone assignments were subsequently aggregated into clone-specific reference profiles, which enabled reference-based assignment of lower-coverage cells. Somatic mutations and copy-number alterations were then aggregated at the clone level to genetically annotate the inferred lineages. Finally, clone identity was integrated with transcriptional state, DNA methylation state, cell-type composition and sampling time point to generate clone-resolved profiles of molecular state, lineage output and longitudinal dynamics.

### Preprocessing and quality control of Droplet-MT data

Raw RNA-seq reads underwent adapter trimming and quality filtering using fastp (v1.0.1). SeekSoul Tools (v1.2.2) was subsequently used to identify and correct cellular barcodes and unique molecular identifiers (UMIs), align them to the human GRCh38 reference genome using STAR (v2.5.2b), and generate raw gene-by-cell UMI count matrices.

Raw DNA methylation sequencing reads underwent adapter trimming and quality filtering using fastp (v1.0.1). The 17-bp cellular barcodes were corrected and appended to the read names using SeekSoulMethyl (v2.1.2). The pooled FASTQ files were then demultiplexed into cell-specific FASTQ files based on the corrected cellular barcodes. Reads from each cell were aligned to the GRCh38 reference genome using Bismark (v0.24.0)^48^, with the maximum insert size set to 1,000 bp (`-X 1000`). The resulting single-cell BAM files were deduplicated using `deduplicate_bismark`, and single-cell DNA methylation calls were generated using `bismark_methylation_extractor`.

For the RNA data, cells with fewer than 300 detected genes or fewer than 400 total UMI counts were excluded. Cells with exceptionally high total UMI counts or numbers of detected genes were also removed as potential doublets. In addition, cells with an unusually high fraction of mitochondrial UMIs were excluded as low-quality cells. For the DNA methylation data, cells were filtered according to the number of covered CpG sites in a task-specific way. In general, cells with fewer than 1 × 10^5^ CpGs were excluded. This cutoff is sufficient for methylation-based cell type embedding and annotation. For de novo lineage reconstruction, cells were further required to have at least 5 × 10^5^ CpGs. For label transfer to shallow query cells, the cutoff was relaxed to just 50,000 CpGs per cell for the T2 cells in AML03.

### Donor demultiplexing of multiplexed libraries

Donor demultiplexing was performed using DNA methylation data at informative SNP-associated CpG sites defined on GRCh38. C>T polymorphisms at CpG sites generate reproducible donor-specific patterns in methylation sequencing, allowing cells from the same donor to be grouped by similarity across many such loci (Extended Data Fig. 2a). For each multiplexed library, a cell-by-locus matrix was constructed at informative SNP-associated CpGs, and cell-cell similarities were calculated and corrected using `methyltree.similarity.compute_similarity_matrix` and `methyltree.similarity.correct_similarity_matrix`. The resulting similarity matrix was clustered to identify donor-specific groups, while cells with ambiguous or mixed-donor profiles were excluded as putative doublets. Donor identities were assigned using separately processed reference cells of known origin by matching each donor-specific group to the corresponding reference population (Extended Data Fig. 2c).

### Transcriptomic preprocessing, integration and clustering

RNA data were analyzed using Scanpy^49^. Each donor was initially processed separately. UMI counts were normalized with `scanpy.pp.normalize_total` and log-transformed with `scanpy.pp.log1p`. Highly variable genes (HVGs) were identified using `scanpy.pp.highly_variable_genes`, followed by principal component analysis (PCA), nearest-neighbor graph construction, Leiden clustering and UMAP embedding using `scanpy.tl.pca`, `scanpy.pp.neighbors`, `scanpy.tl.leiden` and `scanpy.tl.umap`, respectively.

For integrated analyses across donors, cells were combined and donor-associated batch effects were corrected using Harmony^50^. Normalized expression values for HVGs were scaled with `scanpy.pp.scale`, used for PCA, and the resulting PCA representation was corrected with `scanpy.external.pp.harmony_integrate`, using donor identity as the batch covariate. The Harmony-corrected representation was then used to construct the nearest-neighbor graph and generate the integrated UMAP embedding.

### Cell-type annotation

Cell-type annotation was performed using RNA data. Transcriptional clusters from each donor were manually annotated based on the expression of canonical marker genes. In parallel, individual cells were mapped to the BoneMarrowMap reference^51^, and the resulting cell-type labels were used as an independent reference to support the manual cluster annotations. For analyses restricted to CD14^+^ monocytes, we only retained cells with consistent annotation from both manual curation and BoneMarrowMap prediction.

### Doublet detection and removal

For RNA data, Scrublet doublet scores were calculated separately for each donor^52^. Putative doublets were identified from clusters with elevated Scrublet scores and expression of incompatible or mixed cell-type markers. For DNA methylation data, we used cell-type-associated CpGs identified from each donor’s RNA cell type cluster to construct a cell-by-CpG methylation matrix, and cell-cell similarities were calculated with MethylTree, followed by Leiden clustering as described above. Methylation clusters enriched for cells with high RNA-derived Scrublet scores, RNA-defined doublet annotations, or incompatible RNA cell-type identities were classified as doublet-enriched. Cells identified as putative doublets by either modality were excluded from downstream analyses.

### Methylation matrix construction

Single-cell DNA methylation calls were converted into AnnData objects for downstream analysis. Per-cell CpG-level files were loaded with SnapATAC2 using `snapatac2.pp.import_values`, and cell-by-CpG methylation matrices were generated with `snapatac2.pp.make_peak_matrix`^53^. Within each cell, methylation calls from complementary strands corresponding to the same CpG dinucleotide were merged into a single CpG-level measurement. Uncovered CpGs were treated as missing values rather than unmethylated observations.

For supplementary analyses, cell-by-5-kb-bin methylation matrices were also generated by aggregating CpG-level methylation calls across genomic bins. These coarse matrices were used only to evaluate broad methylation structure and unbiased clustering, not for the main epigenetic state embedding or lineage reconstruction.

### Identification of cell-type-associated and lineage-informative CpGs

CpG feature sets were selected from the healthy donors HB10 and F12F. Cells were first grouped by unsupervised clustering of 5-kb-bin methylation profiles, which recapitulated RNA-defined cell types and further separated naive and memory B cells. Cell-type pseudobulks were then scored using `methyltree.lineage.compute_CpGs_variability_across_celltypes` to identify cell-type-associated CpGs and lineage-informative CpGs, the latter characterized by high within-cell-type but low between-cell-type variability. For the identification of lineage-informative CpGs, memory B and T cells were excluded from the cell-type pseudobulk profiles, consistent with their exclusion from lineage analysis, to avoid confounding by the extensive DNA hypomethylation associated with naïve-to-memory differentiation.

Cell-type-associated CpGs were selected independently in HB10 and F12F. The top 2,000,000 loci from each donor were intersected to reduce donor-specific effects, and this shared set was used for cell-state analyses across all donors. Lineage-informative CpGs were selected separately using the same scoring framework. The top 5% of HB10 lineage-informative CpGs (1,402,185 loci) were used for HB10 and all AML donors; F12F reconstruction used its own donor-derived lineage CpG set. Clone-variable CpGs for reference-based assignment were selected separately after reference clones had been established, as described below.

### DNA methylation-based cell-state embedding

Cell-cell methylation similarity matrices enriched for cell-type information were computed from the cell-type-associated CpGs described above. This cell-type similarity matrix underwent further similarity correction, spectral embedding with `methyltree.celltype.spectral_embedding`, Leiden clustering with `methyltree.celltype.leiden_clustering`, and UMAP visualization with `scanpy.tl.umap` (Figure 1h, 2c, 6c, Extended Data Fig. 4d, 5d).

As a comparison, the same similarity-based embedding workflow was applied to 5-kb-bin methylation matrices. This bin-level analysis provided an unbiased view of broad methylation structure but had lower cell-type resolution and was more susceptible to donor-specific variation than the CpG-level cell-type feature set (Extended Data Fig. 1e, 2d, 2e).

### DNA methylation-based cell-type imputation

For cells passing DNA methylation quality control but not RNA filtering, their cell type identities were imputed using cell-cell neighbor graph constructed from the cell-type similarity matrix of the DNA methylation data. For each cell lacking an RNA-derived annotation, we identified its top five closest neighboring cells that have RNA annotation. This cell is then assigned the cell type that has highest total connectivity within this five-cell neighborhood.

### MethylTree lineage reconstruction

For de novo lineage reconstruction, we retained cells with at least 500,000 covered CpGs (approximately 1.7% CpG coverage), while targeting an average coverage of ∼3% or higher across the profiled population. Hypomethylated lymphocytes were excluded to reduce confounding from activation-associated methylation remodeling. Cell–cell lineage similarities were computed using the HB10-derived lineage CpGs for HB10 and AML donors and the F12F-derived lineage CpGs for F12F. Similarity correction, tree reconstruction and clone assignment were performed using `methyltree.analysis.comprehensive_lineage_analysis`.

For datasets dominated by a few clones, similarity correction was recalibrated using inferred clone labels to limit over-correction. Noisy cells were identified with methyltree.qc.identify_noisy_cells. Leiden clusters derived from the lineage-similarity matrix were also compared with inferred clone labels to identify ambiguous or discordant assignments. These ambiguous cells were removed before all downstream analyses, including heatmap visualization, clone-size estimation, clonal-diversity calculations and lineage-bias testing.

### Lineage visualization and summary statistics

Single-cell lineage heatmaps were generated with `methyltree.plotting.plot_lineage_heatmap` using the corrected MethylTree similarity matrix. Clone-level pseudobulk similarity matrices and clone trees were generated with `methyltree.analysis.group_similarity_analysis` and visualized with `methyltree.plotting.plot_tree_with_support`. In the consistency test of clonal assignment (Figure 2i), given clonal assignments from the mixed population, we reported the average accuracy of the reconstructed lineage tree using cells from a purified population against these pre-defined clonal assignments, where the clonal accuracy is calculated using the built-in function `methyltree.metric.compute_lineage_accuracy`. The effective clone number of a population is simply exp(*S*), where *S* is the Shannon entropy of the clone size distribution. This can be computed using `methyltree.clone.clone_diversity_metrics`.

### Reference-based clone-label transfer

We first built donor-specific clonal pseudobulk profiles from the high-depth data. Briefly, de novo lineage reconstruction and clone assignment were performed for high-depth data (above 5 × 10^5^ CpGs per cell). Clones containing more than 20 cells were aggregated into clone-specific pseudobulk profiles. CpGs varying between clonal pseudobulks were selected using `methyltree.lineage.compute_CpGs_variability_across_celltypes`, using clone identity as the comparison variable.

Query cells were compared with reference pseudobulks using Pearson correlations over jointly covered clone-variable CpGs. Correlations were standardized separately for each reference clone across the query population using `scanpy.pp.scale`. Each query cell was assigned to the reference clone with the highest standardized score only if that score exceeded the specified threshold; otherwise, it remained unassigned.

For benchmarking of F12F and HB10 clone-label transfer, cells from library WTJW944 were used to construct reference pseudobulks, whereas cells from library WTJW943 served as queries (both above 500,000 CpGs per cell). For benchmarking of AML03 clone-label transfer using T1 cells, cells with more than 1 × 10⁶ covered CpGs were used to construct reference pseudobulks, whereas cells below this threshold yet passed lineage-reconstruction quality control (500,000 CpGs per cell) served as queries. Query cells were excluded from reference pseudobulk construction and computationally downsampled to the specified CpG coverage, retaining their full-depth clonal assignments as benchmark labels. Queries were classified as reference-represented or reference-absent according to whether their benchmark clone was included in the reference. For reference-represented cells, assignment yield was the fraction receiving a label, and accuracy was the fraction of assigned cells matching their benchmark labels. The false-assignment rate was the fraction of reference-absent cells assigned to any reference clone. Performance across sequencing depths was summarized by the area under the yield–accuracy curve, approximated as the mean product of yield and accuracy across all s.d. cutoffs at each down-sampled coverage level (Fig. 3c,e,f; Extended Data Fig. 3b; Extended Data Fig. 5l).

For AML03 T2, hierarchical assignment accommodated the nested clonal structure. Each query cell first received a tentative label from its most similar reference clone. Query cells were then clustered using `sklearn.cluster.KMeans` on the query-by-reference similarity matrix. A cluster received a clone label when a majority of its cells shared that tentative assignment; ambiguous clusters remained unassigned at this stage. This procedure identified the major compartments and the dominant pClone_1 subclone, pClone1_0. Remaining cells underwent threshold-based assignment to the other reference subclones (Fig. 5l; Extended Data Fig. 5k-m).

### Somatic mutation and copy-number analysis

Candidate AML-associated mutations were identified by clinical targeted sequencing using a 90-gene myeloid malignancy panel. C-to-T (or G-to-A in the complementary strand) substitutions were excluded because they could not be distinguished reliably from conversion-induced sequence changes. The remaining mutations were then queried in Droplet-MT DNA methylation reads using in-house scripts that accounted for nucleotide changes introduced by enzymatic cytosine conversion.

Because single-cell coverage at targeted mutation loci was sparse, mutation evidence was summarized at the clone level. For each mutation, its clone-level mutation fraction was defined as the number of mutation-positive cells divided by the number of cells with callable mutant or reference evidence at that locus. Cells without callable evidence were excluded from the denominator, and absence of a mutant read was not interpreted as a definitive wild-type genotype.

Copy-number profiles were inferred separately from high-depth methylomes (>1 × 10⁶ covered CpGs per cell) using CNVpytor on aligned BAM files with 500-kb bins^54^. A recurrent copy-number signal shared across healthy and AML donors was considered a technical artifact, likely reflecting reduced regional coverage in the methylome libraries, and excluded from downstream analyses (Extended Data Fig. 4h).

### Differential expression and methylation analyses

Differentially expressed genes were identified from log-normalized RNA profiles using `scanpy.tl.rank_genes_groups` with the Wilcoxon rank-sum test. Genes with Benjamini-Hochberg-adjusted *P* < 0.05 and absolute log_2_ fold change > 1 were considered differentially expressed.

Differentially methylated regions were identified with methscan^55^, using the parameter: `methscan diff --bandwidth 2000`, `--stepsize 100`, `--threshold 0.05` and `--min-cells 6`. Candidate DMRs were further required to contain at least 6 CpGs, be covered in at least 20% of cells in each comparison group, have adjusted *P* < 0.05 and show an absolute regional methylation difference > 0.2.

Gene set enrichment analysis was performed with `gseapy.gsea` using the MSigDB Hallmark 2020 gene sets. Gene sets with nominal *P* < 0.05, FDR *q* < 0.05 and absolute normalized enrichment score > 2 were considered significant.

To identify robust malignant-associated molecular differences in CD14^+^ monocytes, differential expression and DMR analyses were repeated across multiple comparison settings: malignant versus recovery-associated monocytes within AML03; AML02 malignant monocytes versus healthy monocytes from HB10 and F12F; and AML01 malignant monocytes versus healthy monocytes from HB10 and F12F. Features that were significantly differentially expressed/methylated across these settings were used to define consensus malignant-associated signatures. In the main text, we illustrated these DMR and DEGs identified from single cells using pseudobulk profiles.

### Pseudobulk similarity to healthy monocytes

CD14+ monocytes were aggregated by donor, ancestry group and time point. RNA pseudobulks retained highly variable genes shared between HB10 and F12F; raw UMI counts were summed within each group, normalized with `scanpy`.`pp.normalize_total` and log-transformed with `scanpy.pp.log1p`. DNA methylation pseudobulks were generated over the cell-type-associated CpGs, not the lineage-informative or reference-clone-variable feature sets.

Pearson correlation matrices were computed separately for RNA and DNA methylation. For each group, similarity to healthy monocytes was the mean of its correlations with the separate HB10 and F12F pseudobulks:

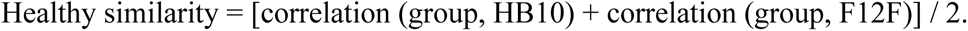

RNA similarities were calculated from expression pseudobulks, not Harmony coordinates.

## Data availability

The Droplet-MT data generated in this study have been submitted to the National Genomics Data Center (https://ngdc.cncb.ac.cn/) under the accession number PRJCA073635. The DARLIN mouse data are available at NCBI GEO: GSE222486.

## Code availability

MethylTree-related code used to reproduce our analysis is available at https://github.com/ShouWenWang-Lab/MethylTree. Additional custom code used in this study is available upon request.

## Acknowledgements

We thank Li Li for her comments, and the Westlake University High-Performance Computing Center for computing resources.

## Author contributions

S.-W.W. and Y.Y.G. conceived the project; J.K.S. performed the data analysis; M.Y.C. carried out the experiments with support from H.Y.L. and M.M.R; Y.L. and J.P. collected the human samples; J.N.S., G.X.Z., Z.-Y.Y. assisted in the development of bioinformatic pipeline; S.S.P. provided feedback on AML data analyses; J.K.S. and M.Y.C. generated figures; S.-W.W. wrote the manuscript with input from J.K.S. and M.Y.C.; S.-W.W. and Y.Y.G. supervised the entire study.

